# Causal modelling reveals lower coral genetic diversity in more heterogenous reef environments

**DOI:** 10.64898/2025.12.16.691338

**Authors:** Ilha Byrne, Iva Popovic, Rebecca H Green, Chancey MacDonald, Jake M Longenecker, Sam J Purkis, Mayra Nunez-Vallecillo, Andrea Rivera Sosa, Antonella Rivera, Pamela Ortega, Katharine E Prata, Kevin R Bairos-Novak, Scott Bachman, Malin Pinsky, Maria Beger, Helen E Fox, Cynthia Riginos

## Abstract

Conserving genetic diversity is crucial for maintaining species’ evolutionary potential and resilience to environmental change. Yet, directly measuring genetic diversity across large spatial scales remains resource-intensive and impractical for conservation planning. Identifying reliable, accessible, and cost-effective spatial predictors of genetic diversity would significantly enhance our ability to incorporate genetic diversity into conservation efforts. Caribbean coral reefs represent a compelling system for investigating such predictors, as these ecosystems face unprecedented threats from disease outbreaks and climate change. Here, we investigate the utility of several environmental proxies as surrogates for genetic diversity, which may be useful for conservation planning in resource- or data-poor regions. To achieve this goal, we used reduced-representation genomic sequencing to estimate intraspecific genetic diversity, and in doing so, identified two genetically distinct and sympatric cryptic coral species in the *Agaricia tenuifolia* species complex. We employed a causal inference approach to assess the relationships between genetic alpha diversity and habitat area and connectivity, habitat heterogeneity, and temperature. Statistical investigations revealed that increased habitat heterogeneity, derived from remotely sensed habitat maps, was associated with lower genetic diversity within coral populations, suggesting that habitat characteristics promoting species diversity may not maintain high genetic diversity within a single species. Habitat area and connectivity, and temperature have predominantly negative effects on genetic diversity with coral habitat area and connectivity showing markedly contrasting responses in two cryptic taxa. Thus, our findings reveal that context matters greatly, as results differ between cryptic species, across variables, and among spatial scales.

## 1 Introduction

Genetic diversity is a fundamental component of biodiversity, providing raw material for adaptive evolutionary change. Higher genetic diversity is associated with greater evolutionary resilience (DeWoody et al. 2021; Kardos et al. 2021), while the loss of genetic diversity is commonly associated with reduced fitness from the expression of recessive deleterious alleles (DeWoody et al. 2021; Kardos and Waples 2024; Shafer and Kardos 2025). Given the central role of genetic variation in supporting adaptive change, the need to safeguard genetic diversity has recently been acknowledged by policy and international benchmarks (Hoban et al. 2021; Hoban et al. 2021). Despite increasing evidence of genetic diversity loss due to anthropogenic activities (Leigh et al. 2019; Shaw et al. 2025), measures of genetic diversity have historically been underrepresented in conservation policy, undermining their relevance to species survival and the maintenance of key ecosystem processes (Bernatchez et al. 2023; Kardos et al. 2021; Laikre et al. 2010). The financial, practical and logistical constraints of assessing genetic diversity across all species and localities of conservation concern (Leigh et al. 2021; Schmidt et al. 2023) motivates the urgent need to investigate scalable approaches to quantify and monitor genetic diversity.

A possible solution for more effectively incorporating genetic diversity into conservation planning is to identify proxies or predictors of genetic diversity that could guide conservation actions (Forester et al. 2022; Hanson et al. 2021; Schmidt et al. 2023). Genetic beta diversity refers to genetic differentiation between populations or sites, whereas genetic alpha diversity refers to the amount of genetic variation within a single population or site. For intraspecific genetic data, beta diversity is often expressed as population-based measures such as isolation-by-distance, isolation-by-environment or genotype-environment associations; such relationships have been extensively documented for many species. These studies suggest that spatial and environmental variables such as temperature and rainfall are often associated with compositional changes in allele frequencies over space and therefore may be useful proxies for genetic beta diversity in terrestrial ecosystems (Wang et al. 2014; Lasky et al. 2023; Rellstab et al. 2015; Selmoni et al. 2021). Proxies for genetic alpha diversity, however, are only starting to be systematically evaluated (e.g., Hanson et al. 2021; Leigh et al. 2021; Schmidt et al. 2023).

Biogeographic theory suggests that landscape characteristics, which encompass both physical habitat features and abiotic conditions, should strongly influence genetic alpha diversity (Bradburd and Ralph, 2019; Kahilainen et al. 2014; Richardson et al. 2014). Habitat area modulates carrying capacity for census population size (*Nc*) and hence genetic drift, where larger habitats can support larger effective population sizes (*Ne*) and therefore greater genetic alpha diversity (Kahilainen et al. 2014; Vellend, 2005). Similarly, greater connectivity among habitat patches results in increased habitat continuity, facilitating gene flow, which can increase genetic alpha diversity. Enhanced connectivity therefore could effectively expand functional habitat area across the metapopulation, collectively promoting the maintenance of genetic diversity within populations (via increased population sizes and reduced genetic drift). In fact, the observation of higher genetic diversity in larger, well-connected populations is entrenched in evolutionary theory (see Hubbell 2011; MacArthur and MacArthur 1961; Vellend 2005) and backed by decades of empirical evidence (e.g., Angelone and Holderegger 2009; Lamy et al. 2013; Schlaepfer et al. 2018). For continuous habitats, however, Neel et al. (2013) suggested that genetic diversity could be elevated under intermediate conditions when overall gene flow is low (allowing genetic differentiation via drift) but gene flow is sufficient to create limited mixing (admixture) between somewhat differentiated genetic groups. Overall, we expect habitat area and connectivity to be intrinsically linked, and we examine these landscape features together throughout this study in relationship to genetic alpha diversity.

Habitat heterogeneity, which we define as variability of habitats or environments within a given area, generates diverse micro-habitats and ecological niches (Kahilainen et al. 2014; Stein et al. 2014). Increased habitat heterogeneity is commonly associated with increased species diversity (Kahilainen et al. 2014; Stein et al. 2014), but its influence on both alpha and beta genetic diversity is unclear (Kahilainen et al. 2014). Specifically, the combined effects of selection arising from environmental differences, the rate of gene flow, and the spatial scale defining a population (relative to the spatial scale of habitat heterogeneity) will modify genetic diversity. For instance, when gene flow is low relative to the spatial scale of environmental differences and sampling area, then genotype-habitat matching could manifest as balancing selection (high genetic diversity) populations arising from local adaptation (Kahilainen et al 2014). Alternatively, when gene flow exceeds the scale of environmental variation, it homogenizes allele frequencies across habitats, preventing local adaptation (Yeaman and Jarvis 2006) and generalist genotypes that can survive under diverse environmental conditions may be favoured, thereby decreasing genetic variation (Kahilainen et al. 2014). Finally, habitat heterogeneity can also regulate population size (Vellend et al. 2005; Kahilainen et al. 2014; Bakker et al. 2024), where increased habitat heterogeneity is expected to reduce the suitable habitat area – and thus depress both *N_c_* and *N_e_* (Kahilainen et al. 2014). In summary, the relationship between habitat heterogeneity and alpha genetic diversity will depend on specific contexts, where characterising the spatial scale of habitat heterogeneity, gene flow, and sampling regime will be important. The two multispecies empirical studies conducted to date found that habitat heterogeneity, characterised by land cover diversity, was negatively correlated with genetic diversity in mammals (Schmidt et al. 2022) but weakly and positively correlated with amphibian genetic diversity (Schmidt et al. 2022), highlighting the often system-specific inferences associated with analyses of genetic diversity. For all the environmental factors discussed thus far, empirical investigations of their effects on genetic diversity are especially lacking in marine environments, providing an opportunity to extend previous terrestrial investigations (Hanson et al. 2017; Schmidt et al. 2022) to marine systems.

While habitat heterogeneity reflects spatial environmental variability, habitats are also variable over time. Temporal environmental variation adds additional complexity to predictions about its impact on genetic diversity. Theory suggests that temporal balancing selection can maintain variation when generation times are short relative to environmental changes (Bergland et al. 2014; Bürger and Gimelfarb 2002; Yeaman and Jarvis 2006). For longer-lived organisms (years to decades), the relevant timescale of temporal variation (whether seasonal fluctuations, interannual variability, or rare extreme events) remains unclear, making predictions uncertain. Moreover, the evolutionary consequences depend not only on the magnitude of variation but also its predictability. Predictable seasonal cycles may favour phenotypic plasticity over genetic diversity, while unpredictable fluctuations may maintain diversity through fluctuating selection (De Jong and Gavrilets 2000; Scheiner 1993). Events that are novel in their extremity, such as unprecedented heatwaves can create population bottlenecks that reduce diversity. Empirical investigations of temporal environmental variation effects on genetic diversity in natural populations remain scarce.

Disentangling multiple factors with direct and indirect effects on genetic diversity is a major challenge of empirical studies (Arif and MacNeil 2023; Schrodt et al. 2025). While observational studies offer advantages such as the ability to span relatively large spatial or temporal scales, it is difficult to make causative conclusions based on observational data (Vellend and Geber 2005). Moreover, confounding variables and measurement errors can create bias and lead to spurious conclusions about causal relationships (Cleemput et al. 2025). One solution is to adopt a causal inference approach, which involves the process of making evidence-based conclusions about the magnitude of a cause-and-effect relationship from observed associations (Cleemput et al. 2025). This methodology has proven valuable in ecological contexts (Arif and MacNeil 2023), such as studying invasive species impacts in South Africa’s Fynbos vegetation (Cleemput et al. 2025) but remains novel in landscape genomics.

In this study, we use a causal inference framework to test whether habitat configuration (area and connectivity), habitat heterogeneity, and temperature influence genome-wide genetic alpha diversity in the critically endangered (IUCN 2020) brooding coral *Agaricia tenuifolia* (Dana 1846) from the Bay Islands of Honduras. Identifying proxies of genetic diversity is particularly relevant to corals as they face unprecedented challenges due to climate change and other local stressors. Coral reefs are also highly heterogeneous and complex environments, characterised by varying levels of connectivity (e.g., isolated reef patches versus contiguous reef networks), environmental heterogeneity, oceanic processes and temperature variability (Bos and Pinsky, 2025; Brown et al. 2023; Longenecker et al. 2025; Pinsky et al. 2023). Moreover, brooding corals, which have short dispersal distances and small effective population sizes, can exhibit genetic structuring over small spatial scales (Prata et al. 2024) perhaps accompanied by local adaptation. In the Bay Islands, coral populations are relatively isolated from other reef systems, minimizing the confounding effects of external gene flow on habitat-genetic diversity relationships. This system and species are therefore well-suited for investigating potential seascape predictors of genetic variation.

Because cryptic species have been described in multiple coral genera (Riginos et al. 2024), including *Agaricia* (Prata et al. 2024) and can differ in environmental responses and genetic variation, we first identify and separate genetically distinct taxa within the *A. tenuifolia* complex to ensure accurate within-population diversity estimates. We subsequently quantify genome-wide alpha diversity for each cryptic species in the context of species-specific dispersal potential and then assess how various environmental variables predict this diversity. By identifying reliable proxies for genetic alpha diversity, our work aims to provide a foundation for rapidly incorporating genetic variation into species distribution models, spatial conservation prioritisation, and restoration planning for coral reefs.

## 2 Methods

### 2.1 Sampling procedure

Sampling locations were selected around Roatán and Utila, the two main islands in the Bay Islands of Honduras, (Western Caribbean; Figure 2D), based on logistical feasibility, and the availability of prior reef monitoring data from the area. Roatán and Utila are compelling case studies because they have experienced severe beaching events and disease outbreaks with some locations historically maintaining high coral cover (e.g., Cordelia Banks, (Keck et al. 2005), and others suffering stark declines in coral cover (Muñiz-Castillo et al. 2019, 2024). Sampling depths did not vary greatly across sites, with all samples collected between 4.2 and 8.1 m. Tissue samples (∼2 cm) of the coral *Agaricia tenuifolia* were collected from 13 locations (26 tidbits) around Roatán and Utila (Figure 2, see Supplementary Materials for additional details on sampling design). Specimens (approximately n = 6 per tidbit) were collected under collection permit Dictamen-ASG-ICF-086-2021 issued by the Forest Conservation Institute of Honduras (ICF). For visualisation purposes paired tidbits were merged i.e., ERA1 and ERA2 into locations i.e., ERA (Figure 2D). Samples were stored in 95% ethanol and shipped to the University of Queensland, Brisbane, Australia, where all laboratory processing was conducted. For population-level analyses, individuals collected at the same tidbit were merged, and only tidbits with appropriate sample numbers (n ≥ 3) were included.

### 2.2 Seascape variables

Environmental variables were assembled from *in-situ* loggers and a suite of satellite products, while the spatial configuration of reef habitat was summarized with a ‘reef-gravity’ index computed for every raster cell in the study area. Our reef-gravity index captures both the amount of reef habitat surrounding each location and its proximity across open water to neighbouring reef habitat patches, providing a biologically meaningful covariate for subsequent analyses (see Supplementary Materials for more details on this metric). Coral polygons from the Allen Coral Atlas (ACA) and and the Khaled bin Sultan Living Oceans Foundation Global Reef Expedition (KSLOF-GRE) / Healthy Reefs Initiative (HRI) (Purkis et al. 2019), together with coastline and land-barrier layers from <u>UN-OCHA</u>, were rasterized at four spatial resolutions (50, 100, 200 and 500 m). Land pixels were masked so that movement could occur only through water. For each focal cell an eight-neighbour SciPy Dijkstra shortest-path search (Dijkstra 1959) was performed through the passable cells out to a distance limit equivalent to 10 hops at the finest resolution (1 km) and to 1-5 km at the coarser grids. Every reachable reef cell 𝑗 contributed 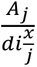 to the focal cell’s score, where 𝐴_*j*_ is the reef area in 𝑗 and 𝑥 is the gravity exponent (variable from 0.5 - 2.5) that weights the influence of distant habitat patches on the focal cell. The resulting rasters of our reef-gravity index thus provide a means of extracting values for each sampling location, with greater reef gravity indicating more contiguous reef habitat.

To estimate habitat heterogeneity, the benthic and geomorphic habitat maps from the ACA database (Lyons et al 2020; Allen Coral Atlas 2022), and the benthic habitat map from KSLOF-GRE/HRI (Purkis et al. 2019) were used. The ACA benthic layer maps six classes (coral/algae, rock, rubble, seagrass, microalgal mats, sand) to ∼10m depth while the geomorphic layer maps 12 classes (reef slope, sheltered reef slope, reef crest, outer reef flat, inner reef flat, terrestrial reef flat, plateau, small reef, patch reefs, back reef slope, shallow lagoon, deep lagoon) to ∼15m depth. Habitat classes in the KSLOF-GRE/HRI map are detailed in the Supplementary Information (Table S1). For each sampling location (tidbit), habitat heterogeneity scores were generated using the Shannon’s diversity index within a predefined spatial window (see Bachman et al. 2023 for details). Window sizes of 1, 5, 10, and 25 Ha were used around each site.

For our reef gravity and habitat heterogeneity metrics, different input maps were compared i.e., Allen Coral Atlas and KSLOF-GRE/HRI maps (see Supplementary Information section 2.4). Moreover, for habitat heterogeneity two diversity indices (Shannon’s and modified Rao’s Q) were compared to determine their influence on resulting measures and statistical inferences (see Supplementary Information section 2.4.2). Given the concordance among maps and metrics, we used the KSLOF-GRE/HRI map supplemented with manual segmentation of live coral patches to estimate reef gravity, and the ACA benthic and geomorphic maps to estimate two measures of habitat heterogeneity.

Between November 2021 and March 2024, 38 temperature loggers (MX2203 HOBO MX Tidbit 400) were installed with 15-minute logging intervals at 18 locations across Utila and Roatán. They were deployed in pairs at a mean depth of 6.2 m ± 1.0 m, with a mean distance between each pair of 295 m ± 94 m. The data corresponding to the years leading up to coral sampling (2021-2023) was used to calculate: mean daily temperature, mean daily temperature range, maximum monthly temperature, standard deviation in monthly temperature, and thermal predictability per site. Thermal predictability was quantified as the product of seasonal amplitude (range of monthly mean temperatures) and the inverse of the coefficient of variation of monthly means. This metric captures both the strength of seasonal patterns and their consistency, with higher values indicating more predictable thermal regimes.

Oceanographic data (mean wave period and significant wave height, and mean wind speed) were also available for certain locations around the Bay Islands. Estimates for each sampling location were approximated based on their proximity to each of five Aqualink buoys (deployed from 2021-2024). For more details on the temperature and oceanographic variables see section 1.4.3 in the Supplementary Information.

### 2.3 DNA extraction, library preparation, and sequencing

Genomic DNA (gDNA) was extracted from a total of 145 samples using the QIAGEN DNeasy Blood and Tissue kit, with an added 30 min RNAse incubation following standard overnight incubation. Quality and yield of gDNA were assessed using gel electrophoresis and extractions were repeated for any low-quality samples. gDNA quantity was then quantified using a Qubit fluorometer (Invitrogen) and normalised to a total 100 ng per sample.

Library preparation was conducted following an in-house double-digestion reduced-representation (ddRAD) approach which employs the enzymes *Pst*I and *Msp*I. The method is adapted from Elshire et al. (2011), Poland et al. (2012) and Petersen et al. (2012), using the Golay adaptor barcodes from Caporaso et al. (2012), (see (Hereward et al. 2020) for more details). This protocol has been successfully applied to corals in the past (Prata et al. 2022) and generates thousands of single nucleotide polymorphisms (SNPs) that can be used to reliably differentiate species and generate genome-wide diversity estimates (Alvarado et al. 2022; Iguchi et al. 2019).

The ddRAD protocol involved enzyme digestion for 3hrs at 37°C followed by 20min at 65°C. Then, an adaptor ligation step was run for 3hrs at 22°C followed by 20min at 65°C. Samples were pooled and size selected for 300-400bp DNA fragments on a Blue Pippin (Sage Science). DNA amplification was performed to ligate sequence indexes, followed by quantification of each library (96 individuals per library). Final libraries were pooled together in equimolar quantities, including a 20% spike-in of a genomic shotgun library. A 0.8x bead-clean was performed to remove any primer dimer present in the final pool. Prepared libraries were sent to the Australian Genome Research Facility (AGRF) for sequencing on an Illumina HiSeq platform (NovaSeq 3000).

### 2.4 Clustering, assembly, variant calling, and initial filtering

Raw sequence data were filtered and assembled *de novo* using iPyrad v0.9.93 (Eaton and Overcast 2020) as there is currently no reference genome for *A. tenuifolia*. Next, we performed stringent data filtering, including contamination removal and clone identification following Bongaerts et al. (2017) and Meziere et al. (2024). Contamination removal was performed using BLASTn comparisons of each locus against six published Symbiodiniaceae genomes (maximum E-value = 10^−15^), and the NCBI non-redundant database to check for non-Cnidarian reads (maximum E-value = 10^−4^). Positive matches were documented and removed from the filtered dataset. Clones were identified and one clonal individual with the least amount of missing data from each pair was retained (19 individuals removed).

### 2.5 Genomic data filtering and population assignment analyses

We performed principal components analysis (PCA; Pearson 1901) using PLINK to identify genetically distinct groups of individuals and assess the effects of missing data thresholds on overall population structure patterns. The dataset used for population assignment analyses was filtered to remove multi-allelic (--max-alleles 2) SNPs, indels (--remove-indels), followed by read depth filtering per locus between 5 and 100 (--min-meanDP --max-meanDP), and a missing data threshold of 50% (--max-missing 0.5) using VCFtools. Next, five samples with greater than 40% missing data were excluded using VCFtools (Danecek et al. 2011). More stringent filtering criteria were then applied to filter sites by minor allele count (--mac 3) and 10% missing data (--max-missing 0.9). Lastly, PLINK v1.9 (Chang et al. 2015) was used to perform linkage pruning (--indep 50 5 2) and to obtain file formats required for population assignment analyses.

To confirm the groupings of genetically distinct individuals, we inspected up to 10 PC axes (Figure S1). To assess population structure and admixture, we used ADMIXTURE (Alexander et al. 2009), with *K* ranging from 2 to 10 genetic groups. Cross-validation errors and likelihoods were calculated for each *K* and used to infer the most likely number of genetic groups. PCA and ADMIXTURE results were visualised using ggplot2 v3.5.2 (Wickham et al. 2016).

The PCA and ADMIXTURE analyses conducted on the entire dataset informed the assignment of individuals into two cryptic taxa co-occurring in sympatry (Figure 2) following guidelines from Riginos et al. (2024) for identifying reproductively isolated species from spatial genetic data. Filtering by read depth, missing data across individuals and number of loci per sample (as above), as well as population structure inference was repeated and conducted separately for each taxon, using the same thresholds and tools as above (Figure 2).

To confirm the nominal species assignment of newly sequenced corals and because *A. agaricites* is also common in the Caribbean and morphologically similar to *A. tenuifolia*, we co-analysed our genomic data alongside genomic data from Prata et al. (2024). We performed a PCA to confirm nominal species groupings and cryptic subgroups to *A. tenuifolia* (see Supplementary Materials section 1.3 for more details).

### 2.6 Isolation-by-distance derived generational dispersal distance

Isolation-by-distance (IBD) is a term used to refer to increased genetic differentiation with geographic distance (Duforet-Frebourg and Slatkin 2016; Wang and Bradburd 2014). Understanding patterns of IBD is crucial to delineating populations and estimating unbiased measures of genetic alpha diversity over a geographic region (Kardos and Waples 2024). Additionally, investigations of IBD enable the estimation of generational dispersal distance, which is an important factor mediating the relevant spatial scale at which the environment is likely to influence intraspecific genetic diversity. To calculate isolation-by-distance (IBD) for each taxon, pairwise genetic distances between individuals were generated using three different genetic differentiation or kinship metrics. Pairwise *F_ST_*, Rousset’s *a* scores (Rousset, 2000), and Loiselle’s kinship coefficient *F* (Loiselle et al. 1995) were calculated using PLINK, genepop v4.7.5 (Raymond and Rousset 1995), and SPAGeDi v1.5. (Hardy and Vekemans 2002), respectively. The slope of linearized genetic differentiation (FST/(1-FST)) versus the log of geographic distance in two dimensions is expected to be proportional to the inverse of Wright’s neighbourhood size, *NS=4πDσ^2^*, where *D* is the effective population density and *σ* measures dispersal as the standard deviation of the probability density describing parental relative to offspring locations (Wright 1943 1946; Rousset 2000; Vekemans and Hardy 2004). As such, slope estimates can be combined with approximated *D* to estimate effective dispersal distances (*σ*). Given that precise *D* estimates were unavailable for our study, we iteratively estimated Wright’s neighbourhood size across a range of *D* values, using estimates derived in a previous study (Prata et al. 2024) on ecologically similar *Agaricia agaricites* (Linnaeus 1758) as a reference.

Because we do not have exact spatial coordinates for each sampled individual, we assigned random coordinates constricted by a 50 m radius around the centroid of each sampling location (for which we have exact coordinates), and a minimum 200 m distance between paired sampling locations (tidbits; see Supplementary Materials section 1.2 for more details). Geographic coordinates were converted to pairwise distances by calculating the Haversine distance between two points. Notably, least-cost pathways (Lewis 2021) were used to account for the physical barriers between sampling locations imposed by land masses when calculating pairwise Haversine distances. Both the assignment of constrained random coordinates to individual coral colonies, and the estimation of pairwise geographic distances were implemented using the packages marmap, geosphere, sp, raster, sf, rasterVis and gdistance within custom python and R scripts.

Genetic and geographic pairwise matrices were compared to generate neighbourhood size (*NS*) and generational dispersal distance (*σ*) estimates. SPAGeDi outputs *NS* and *σ* estimates alongside standard errors by jack-knifing over loci. Genepop outputs Approximate Bootstrap Confidence (ABC) results for the slope and intercept of linearized genetic distance against log geographic distance. A custom R script was used to extract *NS* and σ from the genepop bootstrap results (similar to Prata et al. 2024). Genetic and geographic matrices were used to create a scatter plot of geographic distance on genetic distance (Figure S4). IBD was considered significant when bootstrap 95% confidence intervals did not overlap zero.

### 2.7 Individual- and population-level genetic diversity estimates

To estimate unbiased genome-wide metrics of genetic alpha diversity that account for both invariant and variant sites (see Korunes and Samuk 2021; Schmidt et al. 2021; Sopniewski and Catullo 2024), the demultiplexed and quality filtered FASTQ reads from iPyrad were used as input for Stacks (Catchen et al. 2013). Stacks circumvents the need for a reference genome to generate ‘all sites’ information, whereas iPyrad filters out invariant sites by default. Note that individuals were assigned to one of the two cryptic species and analysed separately in Stacks. Our Stacks workflow followed the recommendations of Sopniewski and Catullo (2024). Loci were constructed *de novo* for each sample (ustacks), setting the maximum distance allowed between stacks in nucleotides (M) to 4. A ‘catalog’ of loci was assembled (cstacks), allowing 4 mismatches (n) between loci. Stacks employs the catalog (record of all the invariant and variant sites in a population) to determine which haplotype alleles are present at every locus in each individual (Catchen et al. 2013). A population map (popmap) was used to call SNPs (gstacks) and generate an ‘all sites’ Variant Call File (VCF). We created a popmap for each taxon, with populations assigned based on tidbit sampling locations.

The Stacks module ‘populations’ was used to generate population-level genetic diversity estimates including pairwise nucleotide difference (*F_ST_*), nucleotide diversity (𝜋), observed heterozygosity (*H_o_*), expected heterozygosity (*H_e_*), and number of private alleles. ‘Populations’ was run iteratively, with a call rate of 1.0 (no missing data) and a mac filter of 1 to remove singletons, including all sequenced individuals and sub-sampling to achieve the same sample size per population. Populations were delineated based on sampling locations given the strong IBD observed in this study system (see Results).

The ‘all sites’ VCF generated for each species was used to estimate individual-level observed heterozygosity (ind *H_o_*) using a custom python script. The VCF was filtered based on best practice recommendations for estimating genome-wide heterozygosity (Hemstrom et al. 2024; Korunes and Samuk 2021). Heterozygosity from single individuals can sometimes deviate from population-wide (pop *H_o_*) estimates, but our individual-level averages closely resemble our population estimates (see Tables S3 and S4). Thus, we proceeded with individual-level observed heterozygosity in our main statistical analyses to maximise sampling numbers, locations and achieve greater statistical power.

Moreover, contemporary effective population size (*N_e_*) was estimated using NeEstimator v2.1 (Do et al. 2014) which interfaces with the R package RLDNe. The bias-corrected version of the linkage disequilibrium method (LDNe; Waples and Do 2008) was implemented with the assumption of random mating (Do et al. 2014). To account for the spatial genetic population but still achieve ample population sizes, we estimated *N_e_* using a sliding window approach in which geographically adjacent sampling locations were iteratively combined to achieve a minimum sample size of 20 individuals. We systematically merged neighbouring locations, then shifted the window along the island (Figure 2D) to create alternative population groupings (Neel et al. 2013). To avoid combining potentially disconnected populations, we constrained the maximum distance between locations within any grouping to the length of Roatán island. This approach yielded three (AT2) to four (AT1) minimally overlapping population groupings per taxon that met sample size requirements and produced successful *N*_e_ estimates.

### 2.8 Causal inference and statistical analyses

Prior to performing statistical analyses, we outlined a set of *a priori* expectations regarding the causal relationships among the variables likely to influence genetic diversity within populations (Table 1). These proposed causal relationships were used to construct directed acyclic graphs (DAGs) that depict the causal pathways among measured and unmeasured (latent) variables (Figure 1). To obtain an unbiased estimate of the causal effect of each focal explanatory variable on genetic diversity, minimally sufficient adjustment sets were determined using ‘adjustmentSets’. In causal inference, minimal adjustment sets are the smallest number of variables to be accounted for (i.e., as covariates) in a model to make a causal effect identifiable by blocking confounding and backdoor paths (Textor et al. 2017). When the variables identified in the minimally sufficient adjustment set are adjusted for (e.g., included as covariates in a regression model), backdoor paths between a treatment and an outcome are blocked, eliminating bias (Textor et al. 2017). The minimal adjustment sets identified in this study are outlined in Table S7. We also conducted a conditional independence check to assess which variables should be independent (unrelated) given certain other variables using ‘impliedConditionalIndependencies’. All of the steps outlined above were conducted using the dagitty package v0.3.4 (Textor et al. 2017). Structural Equation Modelling (SEM) using piecewiseSEM v2.3.1 was conducted to validate conditional independencies and thus the DAG structure. All results associated with the SEM can be found in the Supplementary Materials.

**Figure 1.**
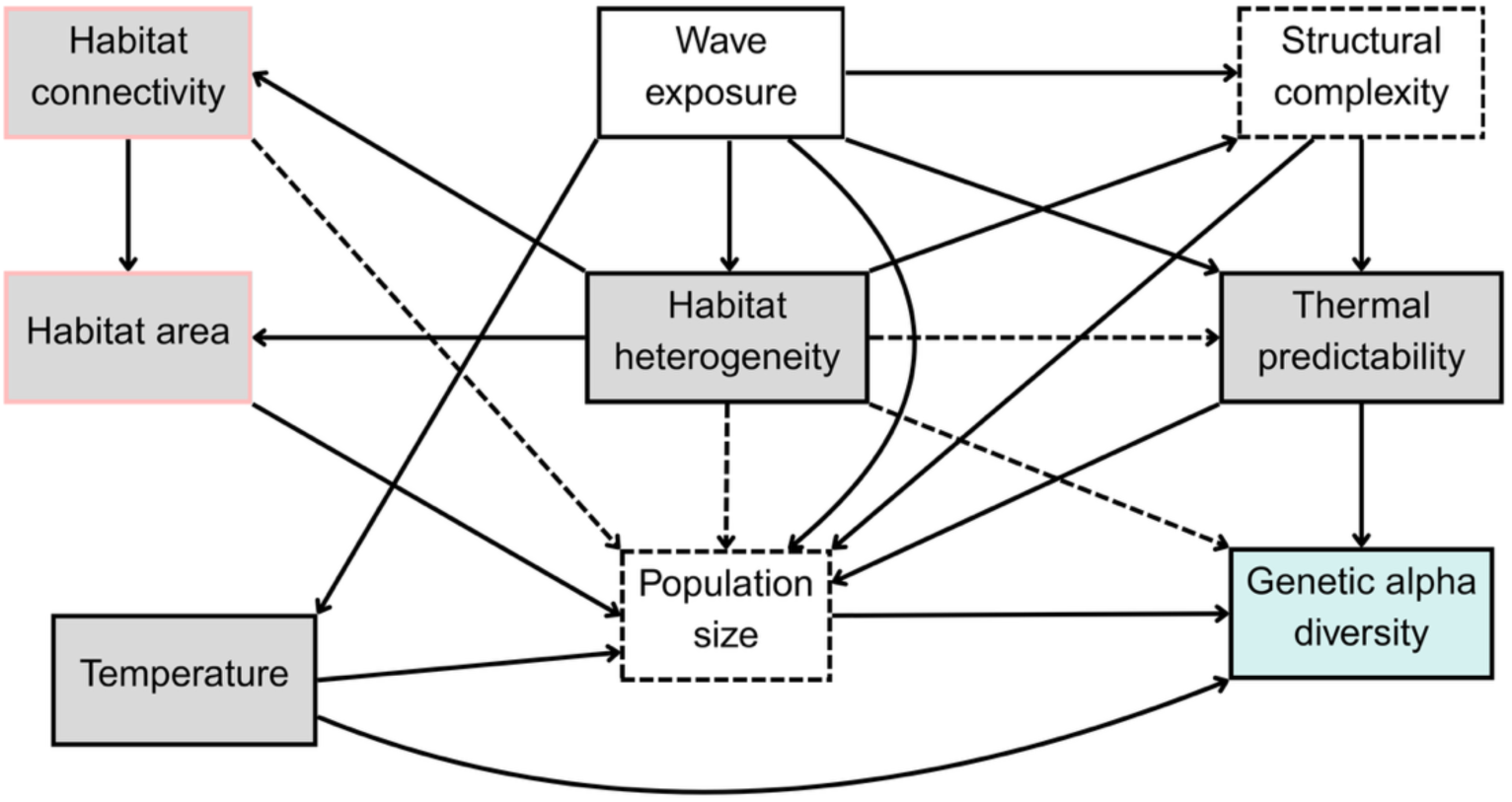
Directed Acyclic Graph (DAG) depicting the relationships among landscape factors that may influence genetic diversity. Arrows are used to depict the predominant direction of an effect between two variables. Ambiguous or uncertain causal pathways are denoted by dashed arrows. Genetic diversity is highlighted as the focal outcome variable in blue. Focal exposure variables are shaded in grey, while habitat area and connectivity are outlined in pink because they are combined into a single ‘reef gravity’ metric in this study. Latent, unmeasured variables with no direct proxies are denoted by dashed boxes. The DAG was constructed based on *a priori* hypotheses (Table 2).

**Table 1.**
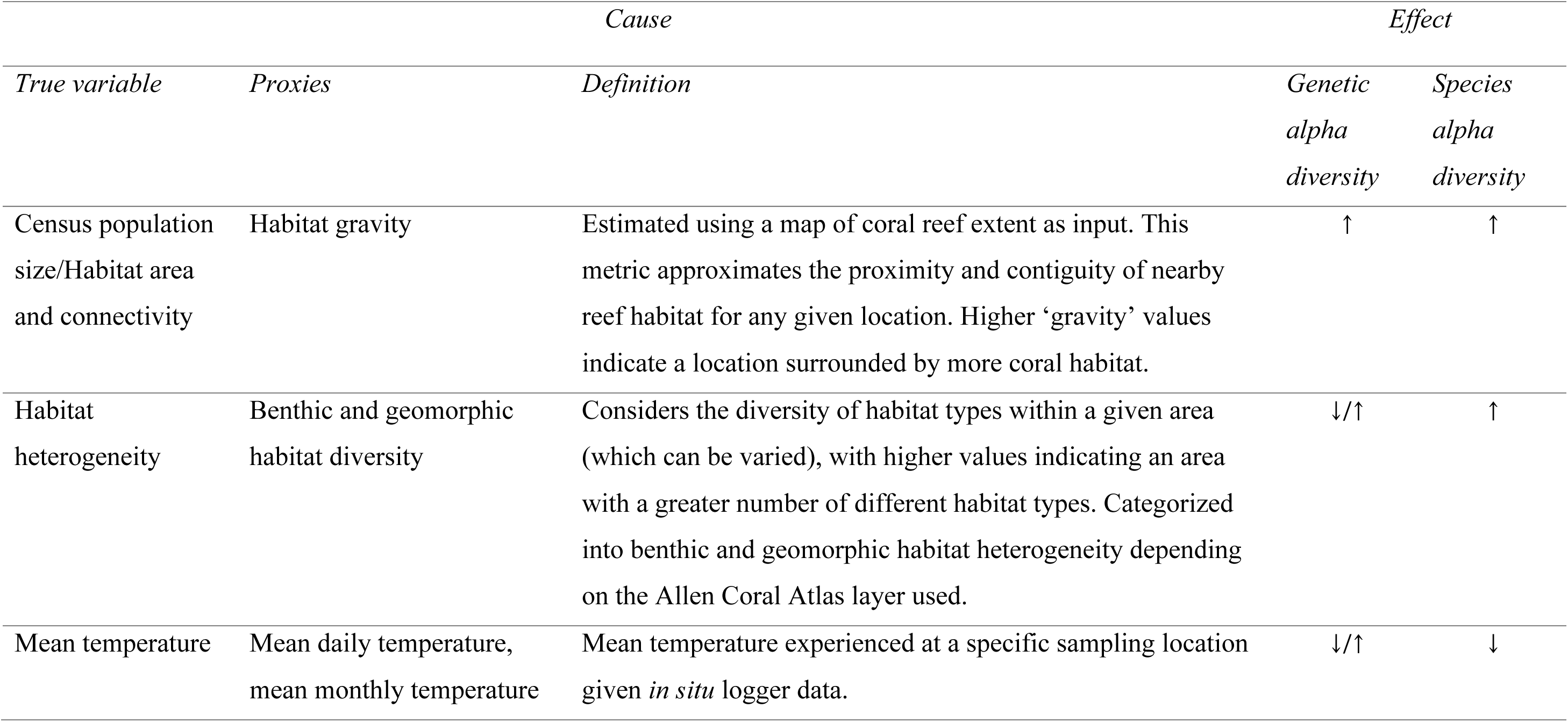
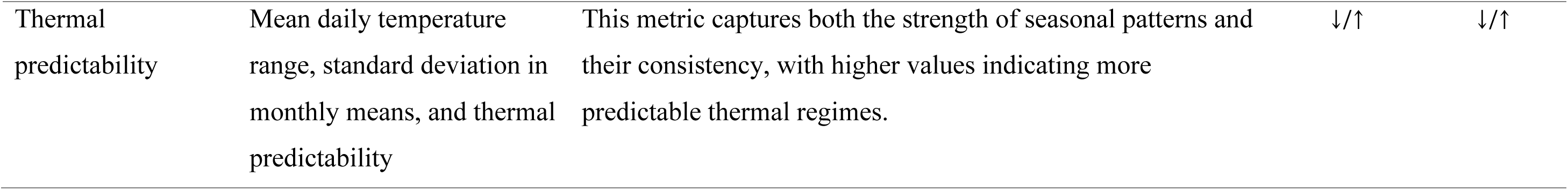
Focal habitat and environmental variables that could modulate levels of genetic alpha diversity. In many cases, the true variable cannot be measured *in situ*, so the proxies used in place of those variables in statistical analyses are listed for transparency. The expected effects of each variable on genetic alpha diversity, genetic beta diversity and species diversity are listed as negative (↓) or positive (↑) based on the theoretical and empirical evidence discussed in the main text.

|  |  | *Cause* |  |  | *Effect* |  |
| --- | --- | --- | --- | --- | --- | --- |
| *True variable* | *Proxies* | *Definition* |  |  | *Genetic*<br>*alpha*<br>*diversity* | *Species*<br>*alpha*<br>*diversity* |
| Census population size/Habitat area and connectivity | Habitat gravity | Estimated using a map of coral reef extent as input. This metric approximates the proximity and contiguity of nearby reef habitat for any given location. Higher ‘gravity’ values indicate a location surrounded by more coral habitat. |  |  | ↑ | ↑ |
| Habitat heterogeneity | Benthic and geomorphic habitat diversity | Considers the diversity of habitat types within a given area (which can be varied), with higher values indicating an area with a greater number of different habitat types. Categorized into benthic and geomorphic habitat heterogeneity depending on the Allen Coral Atlas layer used. |  |  | ↓/↑ | ↑ |
| Mean temperature | Mean daily temperature, mean monthly temperature | Mean temperature experienced at a specific sampling location given *in situ* logger data. |  |  | ↓/↑ | ↓ |

|  |  |  |  |  |
| --- | --- | --- | --- | --- |
| Thermal<br>predictability | Mean daily temperature<br>range, standard deviation in<br>monthly means, and thermal<br>predictability | This metric captures both the strength of seasonal patterns and<br>their consistency, with higher values indicating more<br>predictable thermal regimes. | ↓/↑ | ↓/↑ |

**Table 2.** Causal pathways outlined in the Directed Acyclic Graph (DAG) shown in Figure 1. Bold arrows correspond to causal pathways shown in figure 1, while lighter arrows denote ambiguous/uncertain pathways.

| *Causal pathway* | *Causal link* |
| --- | --- |
| Wave exposure → Structural complexity | Waves and water movement influence erosion, substrate composition and therefore substrate complexity. |
| Wave exposure → Habitat heterogeneity | Waves and water movement modify the physical environment influencing the settlement, community composition and growth of habitat-forming organisms. |
| Wave exposure → Population size | Waves and water movement modify the physical environment by creating disturbance regimes that can influence the growth and survival of habitat-forming and benthic organisms. |
| Wave exposure → Temperature and thermal predictability | Waves and water movement can modulate fine-scale temperature regimes. Exposed reefs are typically cooler than sheltered reefs at the same depth. Areas with higher wave exposure and more water movement are expected to experience less temperature variation. |
| Habitat connectivity → Habitat area | More connected habitat patches effectively increase the total habitat area available. |
| Habitat connectivity → Population size | Habitat connectivity influences the ability of organisms to move or disperse, in turn, affecting gene flow and in turn effective population sizes $N_e$. |
| Habitat heterogeneity → Habitat connectivity | Depending on the scale at which habitat heterogeneity is measured, benthic substrate diversity should strongly influence the area and contiguity of a single substrate type. On coral reefs, higher habitat heterogeneity is expected to result in lower micro-habitat connectivity. |
| Habitat heterogeneity →<br>Structural complexity | The benthic composition of reef habitats (habitat heterogeneity) is expected to influence structural complexity. For example, areas characterised by live coral are expected to be more structurally complex than areas comprised of sand. |
| Habitat heterogeneity →<br>Thermal predictability | The benthic and geomorphic composition of reef habitats (habitat heterogeneity) can influence shading and water movement which will module thermal variation and stress. More heterogenous habitats are expected to be more thermally variable with less predictable thermal regimes. |
| Habitat heterogeneity →<br>Habitat area | In this study, benthic and geomorphic habitat heterogeneity indirectly reflect the amount of live coral within a given area. The benthic composition of reef habitats (habitat heterogeneity) in particular is expected to influence the amount of area comprised of, or suitable for, live corals. Areas with high habitat heterogeneity are expected to harbour less live coral. |
| Habitat heterogeneity →<br>Population size | Given that we expect less live coral in more heterogenous habitats, we can hypothesise smaller effective ($N_e$) and census population sizes ($N_c$) with greater habitat heterogeneity. More heterogeneity is likely to correlate with lower contiguous coral area. However, if gene flow is maintained across microhabitat patches, heterogeneous habitats could maintain large populations composed of generalists with broad niche breadths. |
| Habitat heterogeneity →<br>Genetic alpha diversity | Microhabitat diversity supports local adaptation, possibly increasing overall genetic variation across a metapopulation. Conversely, microhabitat diversity can result in niche differentiation which could result in population differentiation and eventually reproductive isolation, effectively decreasing $N_e$ and also genome-wide genetic diversity. Effects are unclear and dependent on the interplay between selection, gene flow, with sampling scale. |
| Structural complexity → Thermal predictability | Structural complexity creates thermal microhabitats e.g., shaded areas which should affect temperature variation and thermal heterogeneity, which in turn influence the predictability of thermal regimes. |
| Structural complexity → Population size | Increased structural complexity might provide larger surface areas for coral settlement and growth, resulting in larger population sizes. |
| Temperature and thermal predictability → Population size | Temperature can influence the survival and reproduction of corals, with extreme temperature events expected to result in mass coral mortality. Moreover, the variability or predictability of temperatures may influence population sizes, for example, reefs characterised by unpredictable thermal regimes might experience frequent population bottlenecks. |
| Temperature and thermal predictability → Genetic alpha diversity | Temperature in terms of thermal stress is likely to exert a selective pressure on coral populations. The predictability or stability of thermal regimes is also likely to be important. |
| Habitat area → Genetic alpha diversity | Larger populations experience less genetic drift and maintain higher allelic richness. |
| Population size → Genetic alpha diversity | Population size is one of the most important modulators of genetic alpha diversity, with larger population sizes commonly associated with higher genetic alpha diversity (although this relationship is not always positively correlated). |

We used Bayesian generalized linear mixed models to formally test for relationships between genetic diversity and each of our main environmental variables (reef gravity, habitat heterogeneity, mean temperature, and thermal predictability). For habitat heterogeneity, both benthic and geomorphic measures were tested, while for temperature, several different metrics i.e., mean daily temperature and monthly standard deviation were assessed. In all cases, genetic diversity was the response variable, while the predictor variables were determined based on the minimal adjustment sets and thus varied depending on the focal predictor (Table S7).

We fit Bayesian regression models implemented in the brms package v2.22.0 (Bürkner 2017) and further tested the consistency of results across model approaches using spatially-explicit nested Laplace approximation implemented in the INLA package version 22.5.3 (Lindgren and Rue 2015). Unlike Markov Chain Monte Carlo-based methods for fitting Bayesian models, INLA deterministically approximates marginal posterior distributions which allows for faster and more flexible fitting of complex model structures (Beguin et al. 2012). For the INLA models, we accounted for the spatial structure of the data with Stochastic Partial Differential Equations (SPDEs) to remove residual spatial autocorrelation.

Models were first run using all individual genetic diversity estimates (both taxa) with taxon identity included as a fixed effect. Then, because taxa may respond to predictors differentially, we allowed the relationships between predictor and response to vary according to taxa (i.e. using an interaction term resulting in random slopes modelled by taxa). Models were run with both population-level (pop *H_o_*) and individual-level (ind *H_o_*) observed heterozygosity as the response, but we focus on the individual-level results in the Results section as both approaches returned predominantly concordant results (Figure S15). Exploratory models were also run with other population-level metrics of genetic diversity (e.g., nucleotide diversity and expected heterozygosity) yielding similar results to those for *H_o_* (Figure S5). Because our individual-level data were hierarchical i.e., multiple individuals per location, we used a random effect structure to account for variation within localities when modelling ind *H_o_*.

We scaled and centred all variables before analysis so that effect sizes were comparable across models. We used normal priors (normal (0,0.5)) for all parameters in brms. For fixed effects, these were weakly informative priors that do not assume any directionality of effect. Default priors were used for all parameters when modelling with INLA. Model fits were compared using Leave One Out (LOO) cross validation, trace plots, tail effective sample sizes, rhat scores and autocorrelation plots. In cases where the focal predictor variable could be approximated using a number of alternative metrics i.e., in the case for temperature, given that the directionality of effect and model fits were comparable, the model (and metric) with the strongest effect size was selected for reporting.

For habitat heterogeneity, we assessed the influence of spatial extent (i.e., the window size in which habitat heterogeneity was calculated) by running models with habitat heterogeneity estimated at four different window sizes (1, 5, 10, 25 Ha) for both benthic and geomorphic layers. We compared model effect sizes to determine the spatial extent of habitat heterogeneity measurement that best predicted genetic diversity.

Lastly, a ‘kitchen sink’ or ‘causal salad’ model containing all available environmental variables (excluding collinear variables based on Pearson correlation coefficients r > 0.6) as predictors was run in brms to provide a means of comparing the outcomes of predictive versus causal frameworks. All statistical analyses and visualisation described above were performed in R v4.5.0 (R Core Team 2024).

## 3 Results

### 3.1 Genomic data filtering and population assignment analyses

A final set of 122 individuals and 16,722 variant sites (SNPs) were reported post-filtering and used to delineate cryptic taxa. PCA revealed that the sampled *A. tenuifolia* clustered into two genetically distinct, but sympatric groups supporting the presence of two reproductively isolated cryptic taxa (Figure 2A, Figure S9, Figure S10). These genetically distinct taxa occur in sympatry at multiple sites (Figure 2, Figure S9), which suggests that they may comprise two reproductively isolated cryptic taxa within the genus *Agaricia* (Grupstra et al. 2024; Riginos et al. 2024). The PCA results were further supported by the ADMIXTURE results, for which *K* = 2 was the best supported number of clusters based on the highest log-likelihood and lowest CV error (Figure 2E). Taxon-specific PCAs revealed significant geographic population structure in both taxa, which we hereafter refer to as AT1 and AT2 (Figure 2). Here, a superficial comparison of gross morphological features based on in-field colony photographs indicated no obvious differences between taxa (Figure 2). Moreover, co-analysis with *A. agaricites* sequence data (Prata et al. 2024) confirmed that both cryptic taxa (AT1 and AT2) are genetically distinct *A. humilis* species that do not have affinities to other existing species complexes (Figure S8).

**Figure 2.**
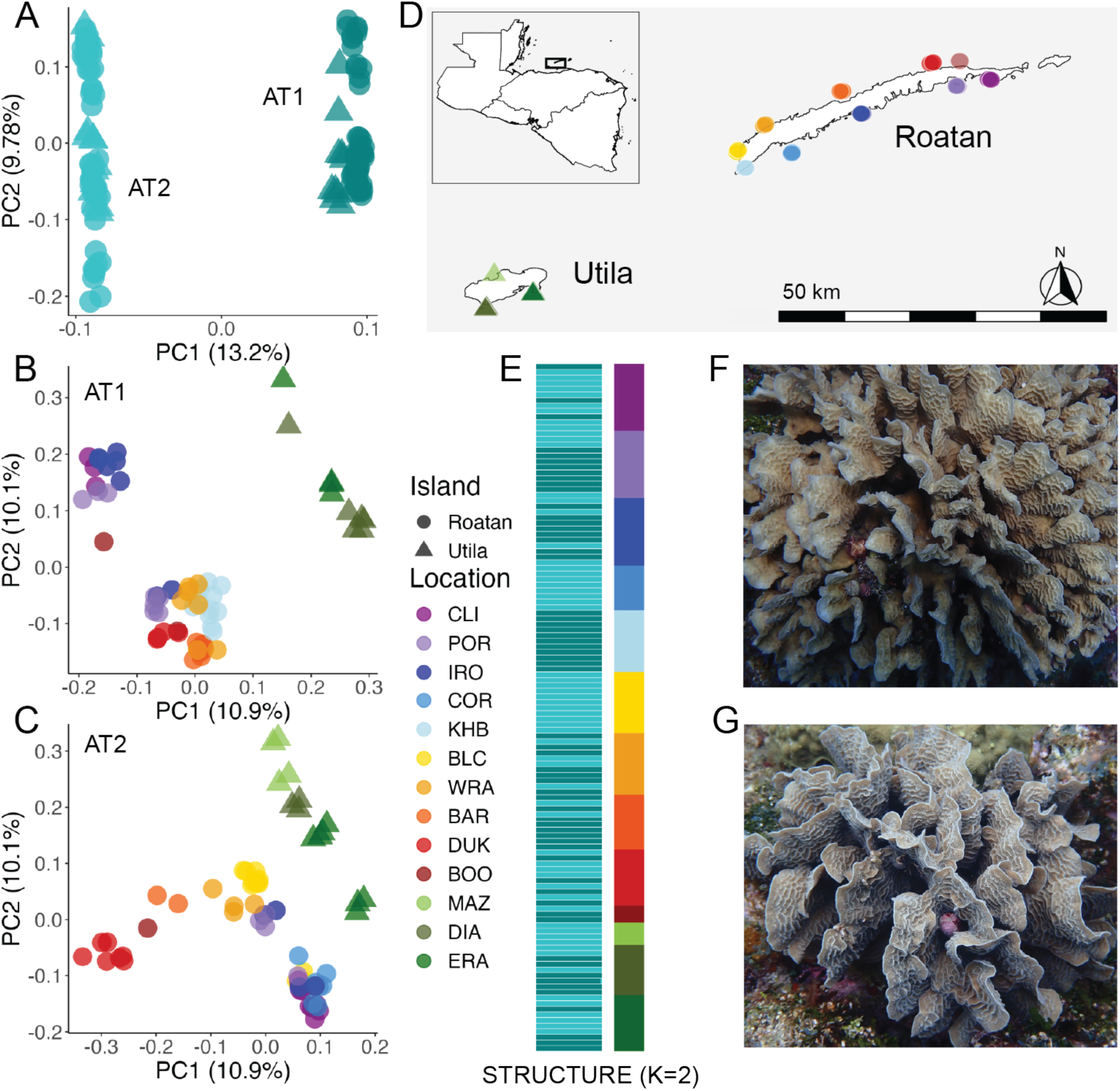
Genotyping reveals two cryptic species in which individual-level variation aligns well with geography. A) PCA plot showing the first two principal components based on ddRAD data from 122 individuals collected in this study. Samples are allocated to one of two clusters, representing two putative cryptic species. B) PCA plot of samples assigned to the AT1 cluster. C) PCA plot of samples assigned to the AT2 cluster. D) Sampling locations around the islands of Roatán and Utila, off the coast of Honduras. E) ADMIXTURE results (with a best supported K=2). Horizontal bars represent individuals, while the colour of each bar represents the putative species assignment (AT1 or AT2). Sampling locations are represented by the colours to the right of the admixture plot. F) A photograph, taken in the field, of a colony assigned to the AT1 cluster. G) A photograph of a colony assigned to the AT2 cluster. Gross morphological differences were not observed between the two species. Field photographs were taken by Pamela Ortega.

### 3.2 Isolation-by-distance derived generational dispersal distance

For both *A. tenuifolia* taxa, significant isolation-by-distance was apparent (Figure S9, Figure S11). The estimates of neighbourhood size were similar between taxa, with 579.70 (± 22.50) estimated for AT1, and 628.80 (± 21.40) for AT2. For both taxa, the median and mean σ estimates across density values were within metres in scale using both Loiselle’s *F* and Rousset’s *â* and regardless of the effective density (*D_e_*) estimate used (Table S5, Table S6). For AT1, the σ estimates ranged from 3–34 m, while the AT2 estimates ranged from 3–35 m, with smaller values reported when larger density values were used due to the relationship between sigma and *D_e_* in the neighbourhood equation (see Table S5, Table S6).

### 3.3 Individual- and population-level genetic diversity estimates

Genome-wide diversity metrics for both taxa are reported in Tables S3 and S4. Overall, genetic diversity was marginally higher in AT2 based on nucleotide diversity (𝜋 = 0.0043, *H_o_* = 0.0024, n = 5) than AT1 (𝜋 = 0.0035, *H_o_* = 0.0024, n = 5). AT2 showed greater variation in 𝜋 (0.0024 to 0.0087) and *H_o_* (0.00097 to 0.0030) among sites compared to AT1 (𝜋: 0.0025 to 0.0047, *H_o_*: 0.0015 to 0.0027). The ERA2 population in AT2 had the highest nucleotide diversity (0.0087), nearly double the highest value in AT1. POR populations tended to have low values of 𝜋 and *H_o_* in both species, while ERA populations tended towards the highest estimates of genetic diversity. Some populations show opposite patterns between species (e.g., BOO1 is high in AT1 but low in AT2, DUK populations show the opposite). Effective population size (*N_e_*) estimates were similar between taxa and ranged between 90 and 210 individuals (Figure 3C). Individual observed heterozygosity was the principal metric used in statistical analyses.

**Figure 3.**
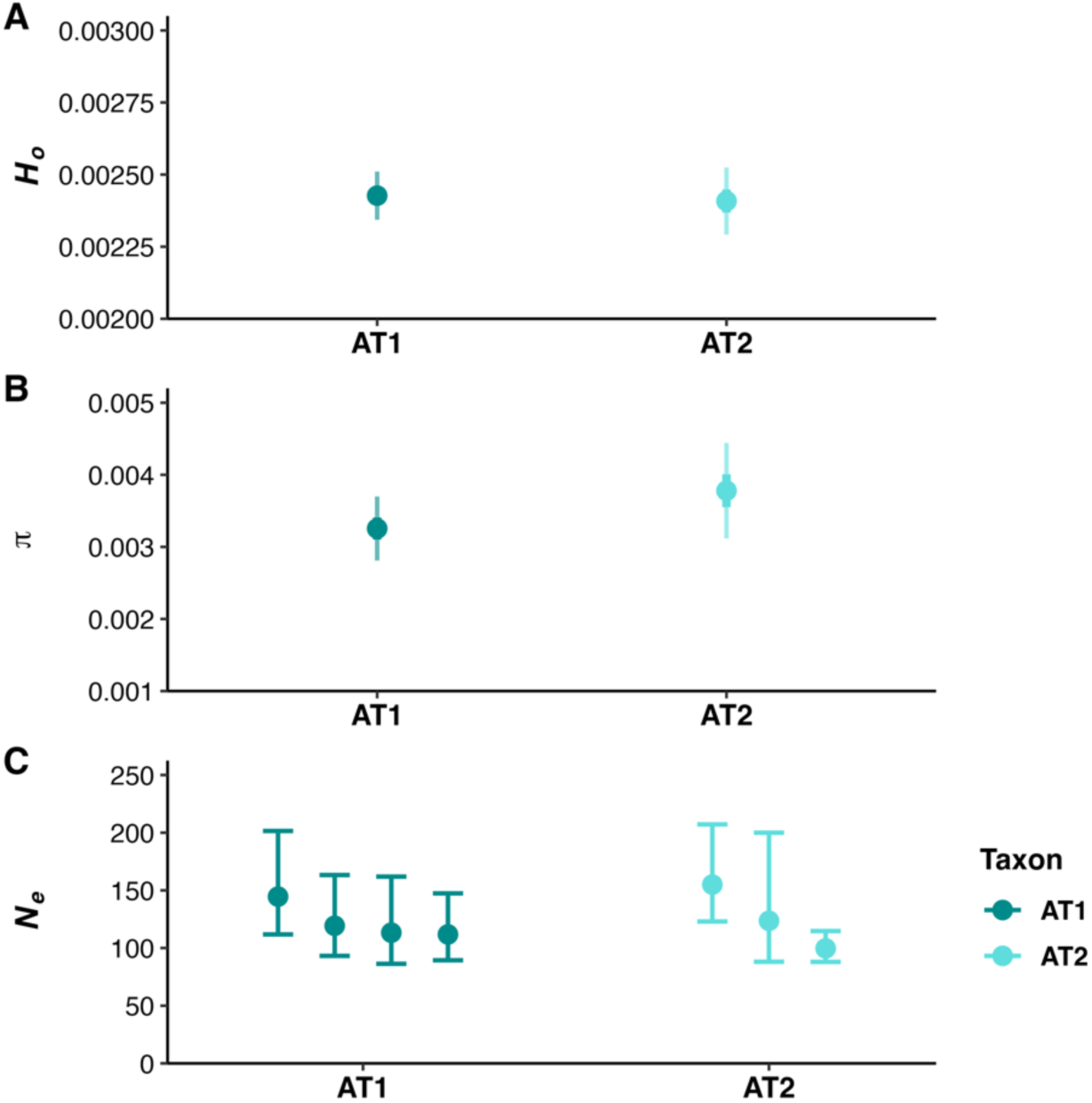
Intraspecific population genetic metrics reveal comparable levels of genetic diversity among two sympatric coral species. A) Estimates of population observed heterozygosity (pop-*H_o_*) calculated using Stacks. B) Estimates of nucleotide diversity (𝜋) calculated using Stacks. C) Estimates of contemporary effective population size (*N_e_*) obtained using the LDNe method within NeEstimator v2.1. Several *N_e_* estimates and their associated parametric confidence intervals are shown, whereby the individuals used to estimate *N_e_* were varied.

**Figure 4.**
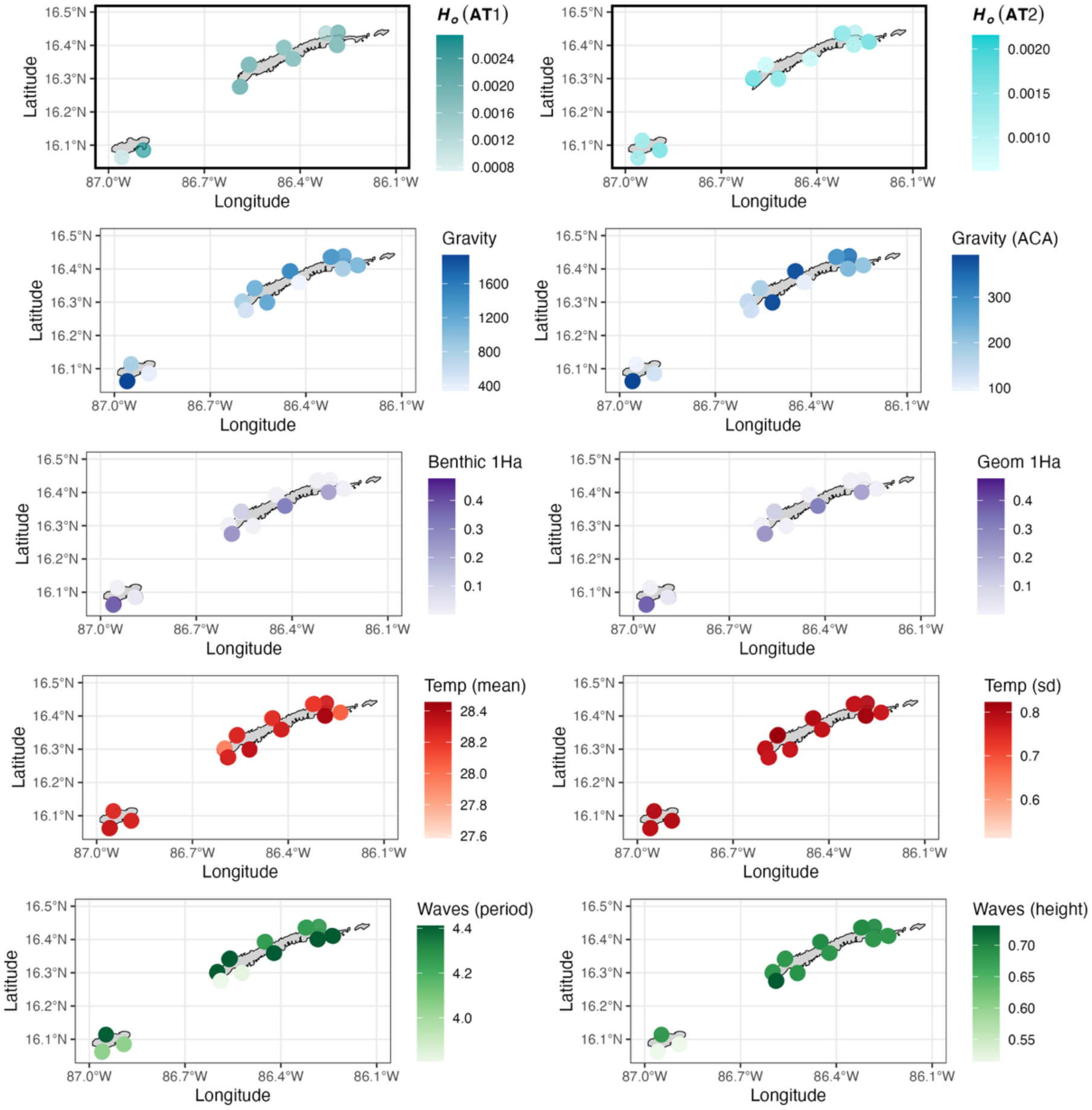
Spatial distribution of genetic diversity (population-level observed heterozygosity) and seascape descriptors across sampling locations in the Bay Islands. Values in green refer to mean individual observed heterozygosity estimates at each sampling location for the two cryptic coral species (AT1 and AT2). ‘Reef gravity’ (Gravity) is shown in blue, values in purple correspond to benthic and geomorphic (Benthic/Geom) habitat heterogeneity (1 Ha spatial extent), and mean daily temperature (Temp) is represented in red.

### 3.4 Seascape variables

Our reef gravity metric showed considerable variation across sampling locations, regardless of which map, or algorithm parameters were used. Locations such as DIA1 and DIA2 (southern Utila) displayed notably elevated values, while sites like MAZ2 (northern Utila) and IRO2 (Roatán) exhibited comparatively lower values, indicating flatter terrain (see Figure 2D for a map of sampling locations). Benthic habitat heterogeneity (1 Ha) ranged from near-zero values at multiple locations to 0.48 at DIA1, while geomorphic habitat heterogeneity was highest at DUK1 (north-eastern Roatán). Wave exposure descriptors such as mean significant wave height and mean wave period exhibited less spatial heterogeneity. The islands are characterized by a low wave energy environment; the significant wave height was below 1 m at all sites for at least 75% of the time. Mean daily temperature displayed limited variation (27.59-28.46°C), though sites CLI1 and CLI2 (north-eastern Roatán) were notably cooler. Monthly temperature variability ranged from 0.51°C at CLI1 to 0.82°C at WRA2, with most sites experiencing similar fluctuations between 0.75-0.81°C, indicating relatively stable thermal regimes across the study region.

### 3.5 Causal inference and statistical analyses

The minimal adjustment sets resulting from our DAG informed our model covariate selection for testing the effect of each exposure variable on genetic diversity (Table S7). Notably, testing the causal effect of thermal predictability on genetic diversity requires blocking a backdoor path through structural complexity (Figure 1: Thermal predictability ← Structural complexity → Population size → Genetic diversity). Because structural complexity is a latent variable we could not directly measure, we excluded thermal predictability from our focal analyses to avoid confounding bias (see exploratory analyses in Figure S16). Despite assessing conditional independencies and DAG support for using SEM, no changes to minimal adjustment sets were required (see Supplementary Information section 2.5).

Bayesian generalized linear mixed models revealed species-specific associations between genetic diversity and our focal seascape variables: reef gravity, benthic and geomorphic habitat heterogeneity and temperature (Figure 5). Without interaction terms (i.e., allowing for species responses to differ), only benthic habitat heterogeneity (1 Ha) showed a detectable effect on genetic diversity (95% CrI excluding zero; Figure 5). However, models that explicitly tested for taxon-specific effects revealed that responses differed in strength between taxa, and in some cases (most notably for reef gravity) in direction (Figure 5). Only the results associated with the effect of gravity on genetic diversity in AT1 and the effects of habitat heterogeneity on genetic diversity in AT2 indicated a 95% credible effect with posterior distributions excluding zero. Results associated with brms models run using population-level genetic diversity metrics and INLA models can be found in section 2.6 in the Supplementary Information.

**Figure 5.**
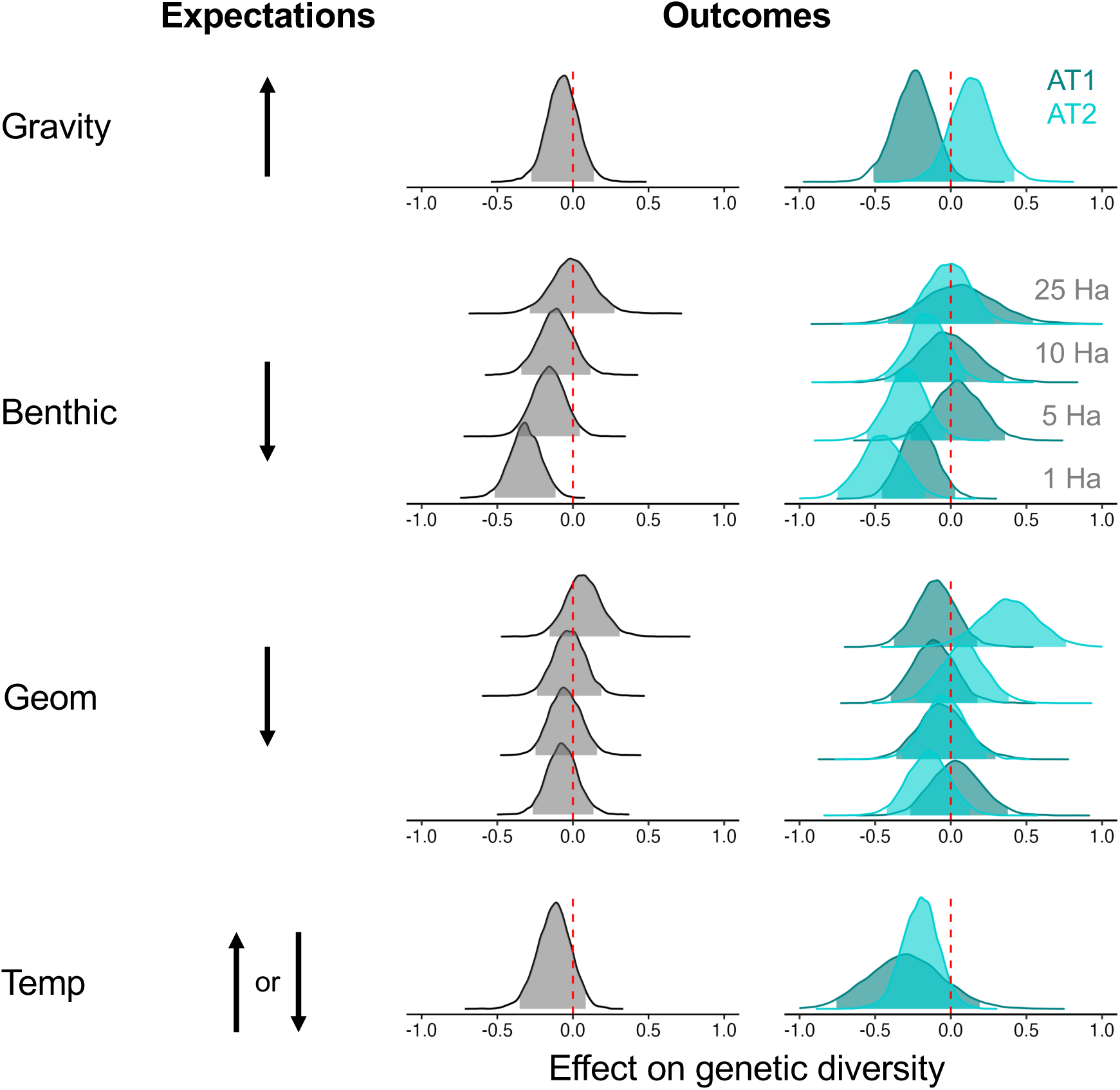
Variable and species-specific variation in the relationship between genetic diversity and focal seascape variables. Rows in the panel correspond to each predictor: reef gravity, benthic habitat heterogeneity, geomorphic habitat heterogeneity, and mean daily temperature. Within ‘benthic’ and ‘geom’, results are shown for four window sizes (25, 10, 5, 1 Ha). Posterior distributions were extracted from Bayesian models assessing the effect of each predictor on overall genetic diversity (both taxa, grey) and species-specific individual-level genetic diversity estimates. Posterior distribution values pertain to standardised posterior values. Credible intervals (95%) are denoted by shaded regions or lines and indicate a 95% probability that the true parameter lies within that range.

For reef gravity, AT1 exhibited a weak negative association with genetic diversity (β = -0.24, 95% CrI: -0.50 to 0.01), while AT2 showed a weak positive association (β = 0.14, 95% CrI: - 0.13 to 0.42). Notably, only the response for AT2 can be considered significant, given the uncertainty associated with the response for AT1 (CrI overlaps zero). The difference between species was substantial, with AT2 showing a significantly more positive response than AT1 (Δβ = 0.39, 95% CrI: 0.03 to 0.74).

Benthic habitat heterogeneity (measured at 1 Ha, the window size most congruent with the spatial extent of sampling at each tidbit) showed a consistent negative relationship with genetic diversity, particularly for AT2 (β = -0.46, 95% CrI: -0.75 to -0.17), for which the posterior distribution excluded zero (Figure 5). Varying the window size (1-25 Ha) across which habitat heterogeneity was calculated revealed varying associations between habitat heterogeneity and genetic diversity. Overall, benthic and geomorphic habitat heterogeneity were mostly negatively associated with genetic diversity across window sizes (Figure 5). For AT1, benthic and geomorphic heterogeneity effects were weak and inconsistent across window sizes, with wide credible intervals indicating substantial uncertainty (Figure 5). However, genetic diversity in AT2 was negatively associated with benthic habitat heterogeneity at 1 Ha and 5 Ha, but positively associated with geomorphic habitat heterogeneity at 25 Ha (β = 0.48, 95% CrI: 0.01 to 0.94).

Temperature effects were similar to those for geomorphic habitat heterogeneity, with both species showing negative but non-credible associations (AT1: β = -0.30, 95% CrI: -0.77 to 0.18; AT2: β = -0.20, 95% CrI: -0.45 to 0.03). The difference between taxa was not credible (Δβ = 0.10, 95% CrI: -0.41 to 0.60), indicating that temperature variation had similar effects on both species.

We also compared effect sizes from the causal models described above with those from a similar predictive model but that included all variables that were non-collinear (Figure 6). While the predictive model achieved slightly higher adjusted R² (0.24 with the predictive model vs. 0.22 with the best causal model), effect sizes differed across variables, particularly for geomorphic habitat heterogeneity (negative in causal, positive in predictive), although differences between models were not significant and credible intervals were wide. The predictive model, by conditioning on gravity, estimates the total effect of geomorphic habitat heterogeneity on genetic diversity (including effects mediated through gravity) which introduces collider bias, producing a putatively spurious positive coefficient.

**Figure 6.**
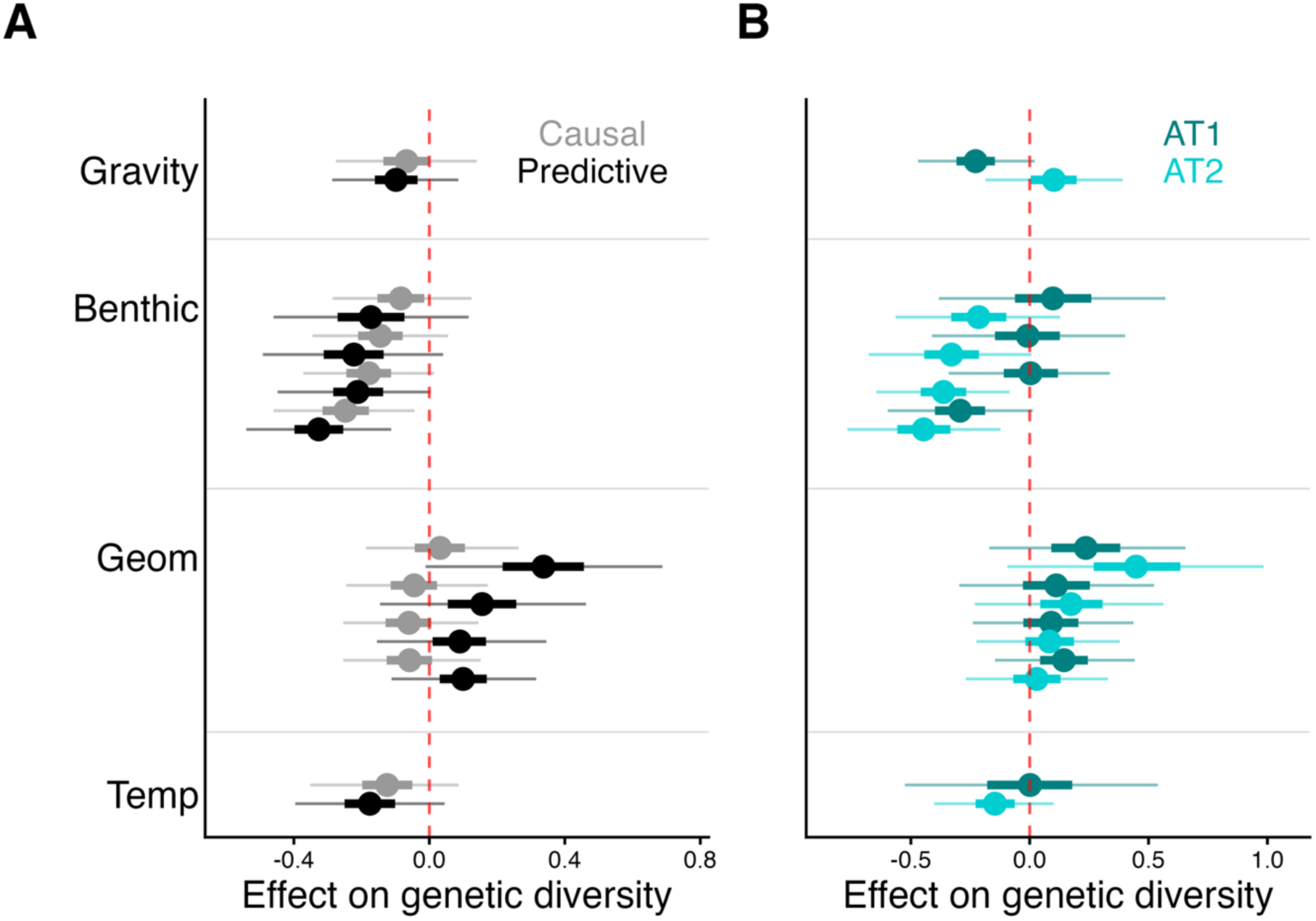
Predictive approaches are not appropriate for drawing causal conclusions. A) Posterior distributions associated with the effect of each seascape variable on genetic diversity extracted from DAG-informed causal and predictive ‘kitchen sink’ models. The distributions for the causal models correspond to those shown in Figure 5. B) Taxon-specific posterior distributions associated with the effect of each seascape variable on genetic diversity extracted from the ‘kitchen sink’ (predictive) model. Within ‘benthic’ (benthic habitat heterogeneity) and ‘geom’ (geomorphic habitat heterogeneity), results are shown for four window sizes in order of decreasing window size (25, 10, 5, 1 Ha). The effect on genetic diversity on the x-axis represents the standardised effect size. The Bayesian R^2^ for the predictive model (containing all variables) was the highest at 0.24, followed by the causal benthic model with an adjusted R^2^ of 0.22.

## 4 Discussion

Identifying causal relationships between environmental variables and intraspecific genetic diversity is critical for understanding the evolutionary processes shaping diversity and forecasting future changes to biodiversity. Using a combination of reduced-representation genomic sequencing, causal inference and Bayesian modelling, we identified a negative effect of benthic habitat heterogeneity on the genetic diversity of two cryptic *Agaricia tenuifolia* species. Our results also implicate the joint effects of decreased habitat contiguity and area on genetic variation. The magnitude and in some cases direction of associations varied when responses were allowed to differ between cryptic species, highlighting the importance of considering discrete genetic groups when investigating environmental drivers of genetic variation. Overall, we examined the potential for environmental proxies to predict genetic alpha diversity in coral reef ecosystems, providing initial steps toward extending previous terrestrial investigations (Hanson et al. 2017; Schmidt et al. 2022) to marine systems.

### 4.1 Two genetically differentiated, sympatrically distributed taxa

A somewhat unanticipated key finding is that *A. tenuifolia* at Roatán and Utila comprises two distinct taxa (Figure 2). This discovery adds to the growing list of nominal species that have been revealed to comprise multiple cryptic coral species with the use of population genomic approaches (Grupstra et al. 2024; Riginos et al. 2024). Despite sympatric occurrence at 10 out of 12 locations, the two taxa were clearly genetically differentiated (Figure 2). Additionally, no indications of admixture between the taxa were observed, with all individuals assigned to a single genetic cluster (Figure 2). The lack of admixture and the presence of distinct genotypes in sympatry are consistent with reproductive isolation due to pre- or post-zygotic mechanisms such as asynchronous reproductive timing (e.g., Chamberland et al. 2025), differential larval settlement preferences, and/or microhabitat niche differentiation. The results that we describe in the following text highlight how environmental predictors and genetic diversity differ between the two cryptic species. Thus, our results reiterate the importance of using genomics to delineate and separately evaluate coral species in ecological studies (Grupstra et al. 2024; Riginos et al. 2024).

### 4.2 Multiple causative factors shape intraspecific patterns of genetic diversity

To our knowledge, this is one of the first applications of casual inference to disentangle the underlying environmental modulators of intraspecific genetic diversity. Causal inference frameworks provide two key advantages for understanding multivariate relationships using observational data. First, they require explicit specification of hypothesised causal pathways, making statistical assumptions transparent and repeatable (Schrodt et al. 2025). Second, they enable identification of direct causal pathways by estimating path coefficients for both direct and indirect effects. Predictive models that ignore causal structure may attribute effects to the wrong variables when predictors are correlated, producing misleading effect sizes that reflect statistical associations rather than biological processes (Arif and MacNeil 2022). We applied these principles to test competing hypotheses about how environmental and habitat characteristics affect genetic diversity in A. *tenuifolia*, focusing on reef gravity, habitat heterogeneity, and temperature.

We hypothesised that increased reef gravity, our proxy for habitat area and connectivity, would support larger population sizes and higher gene flow and thus be associated with higher genetic alpha diversity (Vellend and Geber 2005; Kahilainen et al. 2014; Schmidt et al. 2022a). In line with theoretical predictions, increased reef gravity was positively associated with genetic alpha diversity in AT2 (Figure 5). AT2 exhibits greater spatial population structure than AT1 (Figure 2); thus, at the scale of our study, habitat contiguity may facilitate sufficient gene flow to maintain distinct but connected populations, effectively preserving global genetic diversity (consistent with Neel et al. 2013). For AT1, the relationship between gravity and genetic diversity was negative (Figure 5), contrary to expectations. The results for reef gravity emphasise the importance of distinguishing closely-related species, that respond dissimilarly to environmental stressors, in genotype-environment studies.

Benthic habitat heterogeneity, which incorporates non-coral benthic habitat types from the ACA such as sand and seagrass, effectively captures coral habitat patchiness. In contrast, geomorphic habitat heterogeneity was estimated using reef geomorphic zones (i.e., slope, flat) and therefore the distribution of taxa across zones is likely to determine whether higher geomorphic heterogeneity leads to greater or lesser coral habitat area. Moreover, the spatial scale at which habitat heterogeneity is measured is critical because it determines whether heterogeneity facilitates or constrains selection for habitat matching relative to species’ dispersal capacity (Schiffers et al. 2014). At scales well within dispersal range, habitat heterogeneity may reduce genetic diversity with selection for generalist phenotypes (or matching the most common habitat type), while at scales much larger than dispersal range, heterogeneity may increase genetic diversity by promoting local adaptation and maintaining distinct gene pools with limited connectivity (Yeaman and Jarvis 2006). Given that our estimates of dispersal for both species fall within ∼3-35 m, we expected negative relationships at 1 Ha, but possibly positive relationships at all other spatial scales examined (especially if selection exerts genome-wide effects via linkage; Kahilainen et al. 2014; Vellend and Geber 2005).

Benthic habitat heterogeneity (diversity of benthic habitat types) was negatively correlated with overall genetic diversity (Figure 5). Among the window sizes examined, benthic habitat heterogeneity calculated across the smallest unit area (1 Ha) was the most significant predictor of genetic diversity (Figure 5). However, the finding that habitat heterogeneity is associated with lower genetic alpha diversity corroborates the hypothesis that habitat heterogeneity results in less suitable habitat, effectively reducing *N_c_* and *N_e_* (Kahilainen et al. 2014). Fine-scale habitat features may be important determinants of population demography and genetic diversity in corals with limited dispersal capacity (Prata et al. 2022; Richardson et al. 2014). Schmidt et al. (2022a) similarly found that habitat heterogeneity (land cover diversity) negatively predicted mammalian genetic diversity when measured at dispersal-appropriate scales (5,000–100,000 km²), consistent with our findings for coral benthic habitat diversity.

Our results indicate that the effect of benthic habitat heterogeneity on genetic diversity is more consistent across spatial scales than the effect of geomorphic habitat heterogeneity (Figure 5). The influence of geomorphic heterogeneity on genetic diversity appears to be more scale-dependent, with potentially positive effects emerging at larger spatial scales as seen for AT2 (Figure 5). Thus, benthic habitat heterogeneity at 1 Ha and geomorphic habitat heterogeneity at 25 Ha might be the most useful predictors of genetic diversity in brooding corals as these were the scales at which relationships with genetic diversity were most prominent.

Mean temperature (as well as other metrics of temperature variation and thermal predictability) was the only variable for which neither lumped nor species-specific models detected a significant correlation with genetic diversity. The absence of a strong relationship between temperature and genetic diversity is likely driven by a relatively homogeneous thermal regime across our study system. Moreover, the evidence for a correlation between temperature, or other temporally varying environmental conditions, and genome-wide diversity is generally scarce in the literature (but see Gurgel et al. 2020). Our results align with Thomas et al. (2024) who found no evidence for loss of genetic diversity following periodic disturbances in corals. We conclude that temperature is unlikely to be a useful predictor of genome-wide diversity across small spatial scales, unless habitats are thermally heterogenous. Maximum temperature values (or other statistics that capture extreme thermal events) might be a better predictor than mean temperature, as they are more likely to influence survival and reproduction, and thus genetic variation.

In summary, benthic habitat heterogeneity appears to be the only consistent predictor of genetic alpha diversity, while gravity and temperature variation have weaker direct effects on genetic diversity (Figure 5). Interestingly, the two cryptic species examined here exhibited contrasting genetic alpha diversity relationships with reef gravity despite their morphological similarity, sympatric occurrence, and comparable generational dispersal distances (Figure 2). For cryptic taxon AT2, benthic habitat heterogeneity at 1 Ha and geomorphic habitat heterogeneity at 25 Ha were negatively and positively associated with genetic diversity, respectively. Whereas, for AT1, increased reef gravity resulted in decreased genetic diversity. Patterns for AT2 more closely match our *a priori* expectations (Figure 5). The larger number of sampling locations for AT2 likely contributed to the stronger and more consistent habitat-genetic diversity relationships observed in this species compared to AT1. Furthermore, investigations on selection and local adaptation using outlier analyses and genotype-environment associations could reveal patterns more informative of adaptive genetic variation. Latent spatial patterns or geographic distance likely explain some proportion of the variation in genetic diversity that is not fully captured by the seascape variables explored in this study. Here, the relatively coarse validation of ACA habitat maps (60-90% accuracy), temporally limited environmental data, and low genomic sample numbers, introduce uncertainty when interpreting spatial patterns. Nonetheless, global tools such as the ACA remain useful resources particularly in data poor regions.

### 4.3 Implications for coral reef conservation

Genetic diversity is a fundamental component of biodiversity that has thus far been poorly represented in global efforts to address the biodiversity crisis (Hoban et al. 2021; Laikre et al. 2010). Identifying surrogates for genetic diversity metrics would make it easier to incorporate genetic diversity in conservation frameworks, particularly in resource-limited contexts. Adaptive coral reef management is more timely than ever given the unprecedented heat stress experienced by coral reefs globally (Baker et al. 2025; Muñiz-Castillo et al. 2019). In the Bay Islands of Honduras, *A. tenuifolia* was particularly affected, with field reports of mortality in the majority of all sampled colonies (Rivera-Sosa *pers. comm.*). Here, we outline a robust framework from which surrogates of genetic diversity could be identified. Our findings highlight the importance of incorporating diverse data types and considering evolutionarily significant metrics in spatial conservation prioritisation to ensure all levels of biodiversity are captured and protected.

We also underscore the role of spatial-scale in underlying species-specific responses to seascape heterogeneity, stressing the importance of understanding the life-history and dispersal strategies of focal taxa. Interestingly, our result that benthic habitat heterogeneity is negatively associated with genetic diversity has important implications for spatial prioritisation as conservation frameworks often advocate for the inclusion of diverse habitat types in the selection of protected areas given the established finding of higher species diversity in more heterogenous habitats (Nolan et al. 2021; Bakker et al. 2024). We found that habitat heterogeneity at fine spatial scales was negatively associated with genetic diversity, possibly because fragmented, heterogeneous seascapes reduce effective population sizes through limited gene flow across microhabitats. In contrast, at broader landscape scales, habitat heterogeneity may indeed enhance regional genetic diversity by maintaining differentiated populations across environmental gradients, as predicted by theory. Our results therefore highlight that local habitat diversity may constrain genetic variation within populations, possibly due to strong local selection pressures or restricted gene flow across heterogeneous microhabitats, even while broader-scale habitat heterogeneity maintains regional genetic diversity through population structure. This scale-dependency highlights the need to consider both local and regional processes when considering habitat diversity as a surrogate for genetic diversity in conservation planning.

Metrics that capture elements of habitat area and connectivity, such as our reef gravity metric, could be promising predictors of genome-wide genetic diversity given fundamental principles in ecology and evolutionary biology (Kahilainen et al. 2014; Vellend 2005; Vellend and Geber 2005; Wang and Bradburd 2014). Most landscape genetics and conservation genetic recommendations would argue that protecting larger and more contiguous patches of habitat would support the evolutionary resilience of most species (Forester et al. 2022; Schmidt et al. 2024). However, we demonstrate that the relationship between habitat configuration and genetic diversity is more nuanced than the simple assumption that larger, more connected habitats support greater diversity. Genetic differentiation, and the protection of unique or locally adapted genotypes, should not be overlooked (Bernatchez et al. 2023) to ensure rare genotypes are represented in management plans (Nielsen et al. 2023; Riginos and Beger 2022). Linking genomic variation to fitness outcomes, thermal tolerance, and adaptive capacity is urgently needed to inform when and where genetic diversity should be incorporated in conservation decision-making.

Furthermore, our estimates of dispersal and *N_e_* corroborate previous studies that have found restricted, metre-scale dispersal among brooding coral species (Prata et al. 2024). This highlights the potential vulnerability of brooding coral species, that mostly rely on local replenishment, to future environmental disturbances. Understanding these species-specific dispersal and demographic parameters is critical to designing effective spatial management strategies. Overall, our results underscore the importance of considering habitat contiguity and configuration in protected area design, especially for brooding coral species with limited dispersal capacity.

## Acknowledgements

The authors thank all CORAL team members who supported and participated in field collections. Collections of coral specimens were conducted under research permit Dictamen-ASG-ICF-086-2021. Data storage, computing and bioinformatics were facilitated by the University of Queensland (UoQ) agreement with the Queensland Cyber Infrastructure Facility. We thank members of the Riginos Research Group for reviewing and supporting various aspects of the work contained within this manuscript. We also thank members of the CORAL research team and collaborators who provided insightful comments throughout the project.

## Data Availability Statement

Data and scripts associated with this study will be made available upon publication in a peer-reviewed journal.

## Generative AI Statement

Generative AI tools were not used in the writing of this manuscript. However, AI tools were used to aid literature searches e.g., Connected papers, and to troubleshoot and debug errors in R code e.g., Claude.

## Supporting Information

See attached Supplementary Information document for additional figures, and details relating to methods and results.

