## Supplementary information for "Causal modelling reveals lower coral genetic diversity in more heterogenous reef environments"

**Table of Contents**


| | |
|---|---|
| **1.0 Materials and methods** | **3** |
| 1.1 Sampling procedure | 3 |
| Figure S1 | 3 |
| 1.2 Caribbean *Agaricia spp.* delineation | 4 |
| 1.3 Environmental data sources | 5 |
| 1.3.1 Allen Coral Atlas | 5 |
| Figure S2 | 5 |
| Figure S3 | 5 |
| 1.3.2 Healthy Reefs Initiative | 6 |
| Figure S4 | 6 |
| Table S1 | 6 |
| 1.3.3 Custom reef extent maps | 6 |
| 1.4 Environmental variables | 8 |
| 1.4.1 Reef gravity | 8 |
| Figure S5 | 8 |
| 1.4.2 Habitat heterogeneity | 9 |
| Figure S6 | 9 |
| 1.4.3 Temperature metrics | 10 |
| Table S2 | 10 |
| 1.4.4 Autocorrelation amongst variables | 11 |
| Figure S6 | 11 |
| 1.5 Structural Equation Modelling | 12 |
| **2.0 Results** | **13** |
| 2.1 Caribbean *Agaricia spp.* delineation | 13 |
| Figure S8 | 13 |
| 2.2 *A. tenuifolia* cryptic lineages | 14 |
| Figure S9 | 14 |
| Figure S10 | 15 |
| 2.3 Intraspecific analyses | 16 |
| Table S3 | 16 |

Table S4 16
Figure S11 18
Table S5 19
Table S6 20
2.4 Environmental variable optimisation 21
2.4.1 Reef gravity 21
Figure S12 21
2.4.2 Habitat heterogeneity 21
Figure S13 21
2.5 Causal framework 22
2.5.1 Minimum adjustment sets 22
Table S7 22
2.5.2 Structural Equation Modelling 23
Table S8 24
Table S9 24
Table S10 25
Figure S14 25
2.6 Model comparisons 26
2.6.1 Individual versus population responses 26
Figure S15 26
2.6.2 Effect of temperature on genetic diversity 27
Figure S16 27
2.6.3 Spatially explicit models 28
Figure S17 28
2.7 Bayesian GLMMs extended tables 29
Table S11 29
Table S12 29
Table S13 30
**3.0 References** **31**

**Materials and methods**

1.1 Sampling procedure

Coral colonies were sampled at 13 locations around the Bay Islands as shown in the main text (Figure 2). However, each location e.g., POR consisted of paired sampling tidbits (e.g., POR1 & POR2) situated approximately 200 m apart. At each tidbit, six coral colonies were sampled in three radial transects of 15 m (two samples per transect, each 10 m apart). Colonies were only sampled if they were at least 10 cm in diameter. A total 145 samples were collected, 28 from Utila and 117 from Roatán. Coral tissue was placed in 95% ethanol for long-term storage. The ethanol was changed three to four times, upon placing tissue in ethanol, 1-4 hours after the initial change and then another change in ~24 hours.

For visualisation purposes paired tidbits were merged i.e., ERA1 and ERA2 into ERA i.e., in Fig. S9. For population-level analyses, only tidbits with appropriate sample numbers (n ≥ 3) were included (due to DNA extraction and sequencing failures not all samples resulted in quality sequence data).

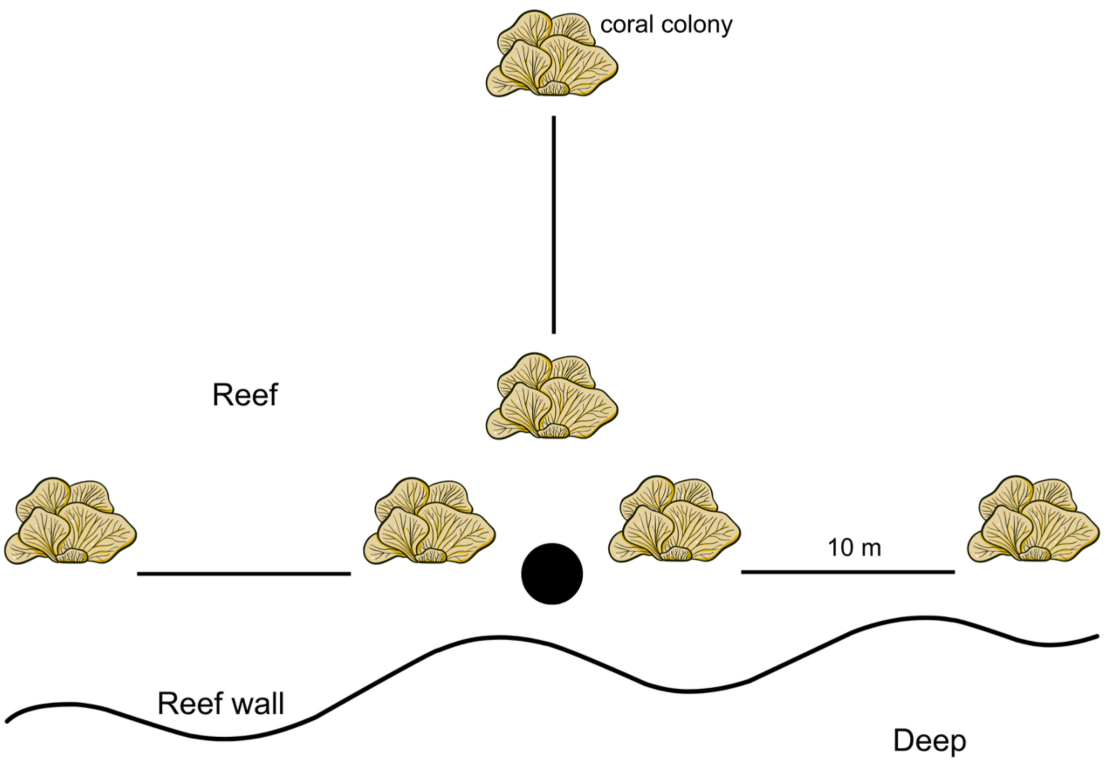


**Figure S1** A schematic representation of the tidbit sampling design adopted in this study. At each tidbit location, three radial transects were surveyed and two colonies, 10 m apart, were sampled per transect.

1.2 Caribbean *Agaricia spp.* delineation

To assess the possibility that one of the Agaricia tenuifolia taxa identified in this study could belong to the nominal species *A. agaricites*, a subset of our data was analysed alongside sequence data (associated with *A. agaricites* and *A. humilis* sampled in Curaçao) previously generated by Prata et al. (2024). We selected 10 individuals per cryptic lineage (AT1, AT2, AA1, AA2) as well as 10 *A. humilis* individuals and performed denovo assembly and variant calling using ipyrad v0.9.105 with default parameters. Further quality filtering was performed on the VCF file generated using ipyrad, including the removal of indels, applying a minimum and maximum allele threshold of two, and only retaining SNPs that were present in >50% of individuals using VCFtools v0.1.16. Additional filtering for biallelic sites with a minimum allele count of three was conducted and reads with low (<5) and high depths (>500) were removed. Finally, linkage pruning was conducted by applying an $R^2$ threshold of 0.2 using PLINK v1.9. PCA was conducted using PLINK and visualised in R using ggplot2. *A. agaricites* and *A. humilis* individuals were coloured based on their cryptic lineage assignments as previously determined (Prata et al. 2024).

### 1.3 Environmental data sources

#### *1.3.1 Allen Coral Atlas*

Benthic and geomorphic habitat maps were downloaded from the globally available Allen Coral Atlas (ACA) database, which maps the world's shallow water reefs with a spatial resolution of 5 x 5m (Lyons et al 2020, Allen Coral Atlas 2022).

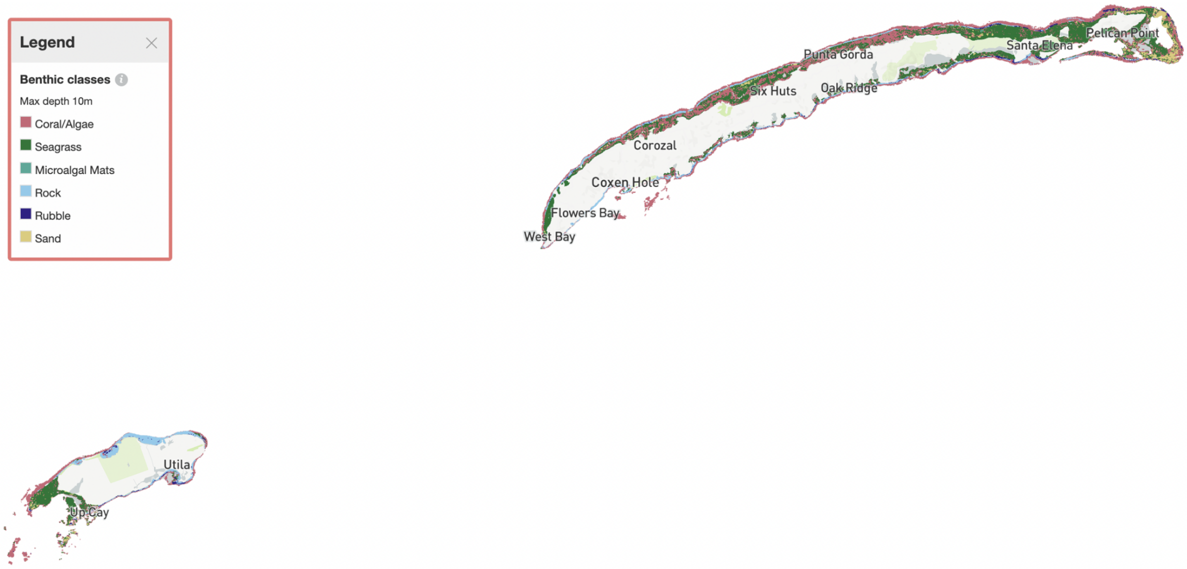


**Figure S2** The Allen Coral Atlas (ACA) benthic habitat map used to estimate and extract benthic habitat heterogeneity in this study. Obtained from the ACA public website (Lyons et al. 2024).

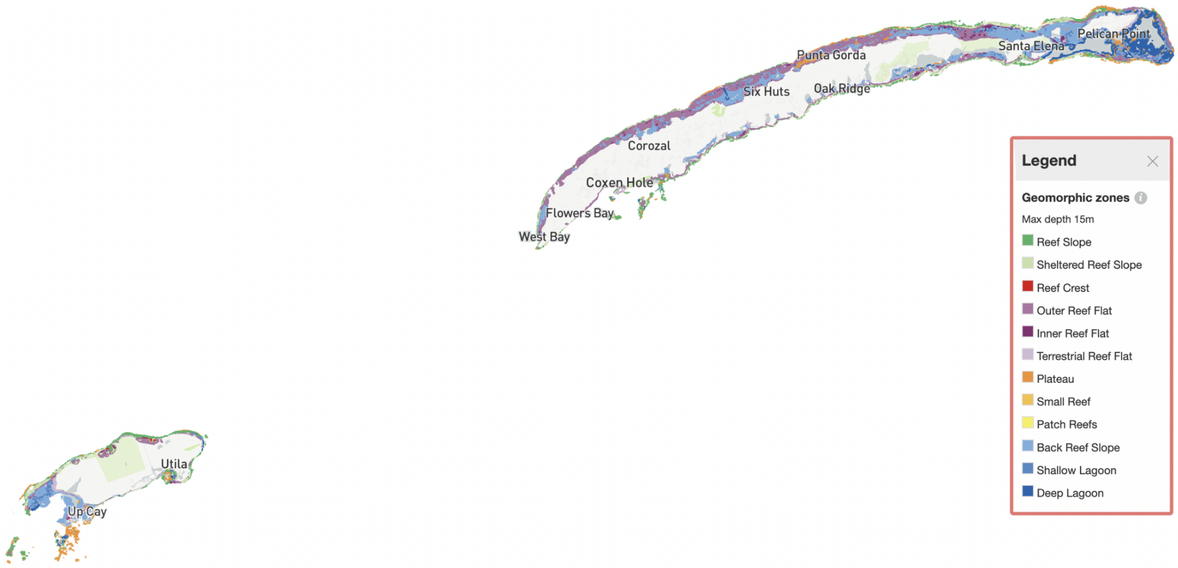


**Figure S3** The Allen Coral Atlas (ACA) geomorphic zone map used to estimate and extract geomorphic heterogeneity in this study. Obtained from the ACA public website (Lyons et al. 2024).

*1.3.2 Healthy Reefs Initiative*

Additional coral reef habitat maps were assembled using information from AGGRA and the Healthy Reefs Initiative. In Honduras, the creation of these maps was supplemented with expert insights to improve their accuracy by removing erroneous classifications. These maps and associated resources are available in Purkis et al. (2019).

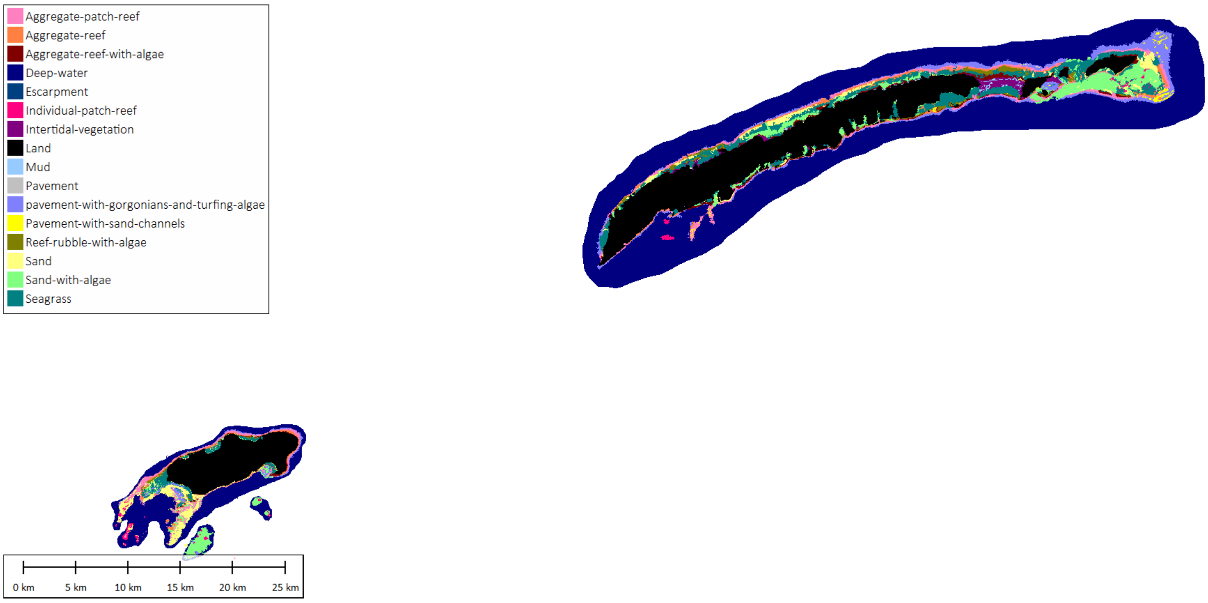


**Figure S4** The Healthy Reefs Initiative (HRI) habitat map used in this study to estimate and extract habitat variables such as habitat heterogeneity and reef gravity.

**Table S1** Descriptions of habitat classes used in Fig S4. Modified from Purkis et al. (2019).

| *Habitat class* | *Description and notes* |
|---|---|
| Aggregate reef | Continuous reefs where individual patches exceed 250m in diameter. |
| Aggregate reef with algae | Areas that are too shallow or inclement to sustain coral communities based on the colouration of satellite imagery. |
| Aggregate patch reef | Individual patch reefs that are aggregated because they are smaller than 250m in diameter and due to image resolution are unable to be discretised as individual entities. |

| | |
|---|---|
| Individual patch reef | Isolated coral reef formations that are circular or elongate in shape. Often surrounded by a highly reflective ring of bare sand. |
| Reef rubble with algae | Grey-to-green-to-brown colouration and texturally homogenous in satellite imagery. |
| Seagrass | Dark green to black with sinuous texture in imagery. |
| Sand with algae | Tinged grey in satellite imagery as opposed to the highly reflective sand class. |
| Sand | Areas that appear highly reflective and homogenous in satellite imagery. |
| Pavement with gorgonians and turfing algae | Hard bottom covered in a thin sheet of sand, or sparsely colonised by turf algae, sponges and rare coral colonies. Often dark blue to black and smooth in texture in imagery. |
| Pavement with sand channels | Hard bottom covered with unconsolidated sheets of sand. Visible as bright beige to white in imagery. |
| Vegetation | Includes mangroves, palm trees and coastal shrubbery. Appears as bright green in true-colour satellite imagery. |
| Mud | Fine-grained sediments found at 0-2m depths. Appears light grey to dark brown in imagery. |
| Land | All terrestrial habitats whether man-made or natural. Characterised by a high infrared spectral response. |

*1.3.2 Custom reef extent maps*

Custom reef extent maps were created using ArcGIS by manually outlining reef polygons using a google earth base layer. Polygon positioning was supplemented with knowledge of known coral reef areas and information from the HRI map. Polygons were exported as a raster file and used to create our reef gravity index.

### 1.4 Environmental variables

#### *1.4.1 Reef gravity*

The raster classification treats any overlap with land as an impassable land pixel. There are trade-offs to this detection, but it is appropriate for our study system due to the high spatial heterogeneity of land and density of small islands in the study region.

Each water pixel's gravity score is the sum of influences from all coral pixels that can be reached by traveling through water (land blocks paths) up to a set maximum range. Every coral pixel contributes equally, and its influence fades with increasing water-path distance based on a decay exponent i.e., $gravity_i = \sum \frac{pixel\ weight^{\frac{1}{pixel\ distance}}}{number\ of\ pixels}$. Distances are measured on an eight-neighbour grid (cardinals and diagonals), self-influence is excluded, and unreachable sources do not contribute. The three main parameters are map raster resolution, the maximum flow distance, and the decay exponent, which together control how fast influence decays and how far it spreads. The 'weight' is measured in spatial degrees of contribution, and the gravity map is a relative index with units of inverse degrees to whatever decay is chosen.

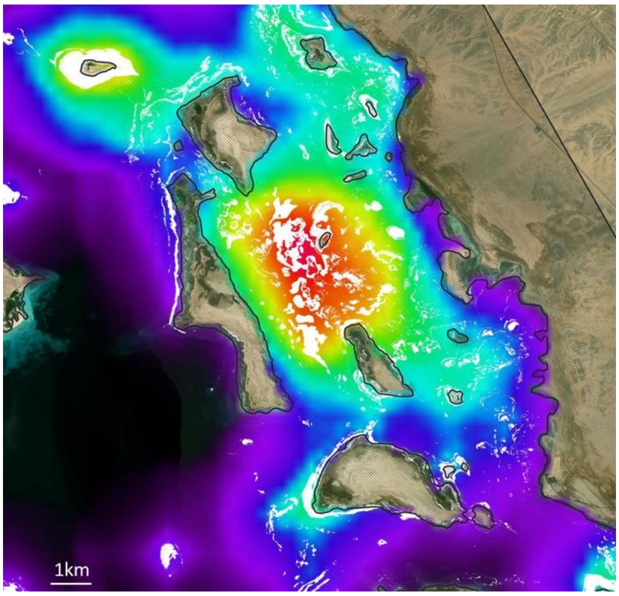


**Figure S5** An example representation of the raster output associated with the reef gravity index. The white areas symbolise mapped coral habitat, the brown thatched areas symbolise land. The red areas are identified as areas with highest reef gravity while the purple areas are the lowest gravity regions.

*1.4.2 Habitat heterogeneity*

For each location, habitat heterogeneity scores were generated using Rao's Q metric, which is normalized to account for the presence of any null values (land or deep water) within the window index (see Bachman et al. 2023) for details) and the Shannon's diversity index (see Bakker et al. 2024 for details). We refer to Rao's Q as 'modified' because it was weighted based on the percentage of valid points in the window. This was to prevent unrealistically large Q values from appearing where there were barely any valid points. Window sizes of 1, 5, 10, and 25 Ha were used around each site. The final set of habitat heterogeneity values used in statistical analyses were generated using Rao's Q (Bachman et al. 2023).

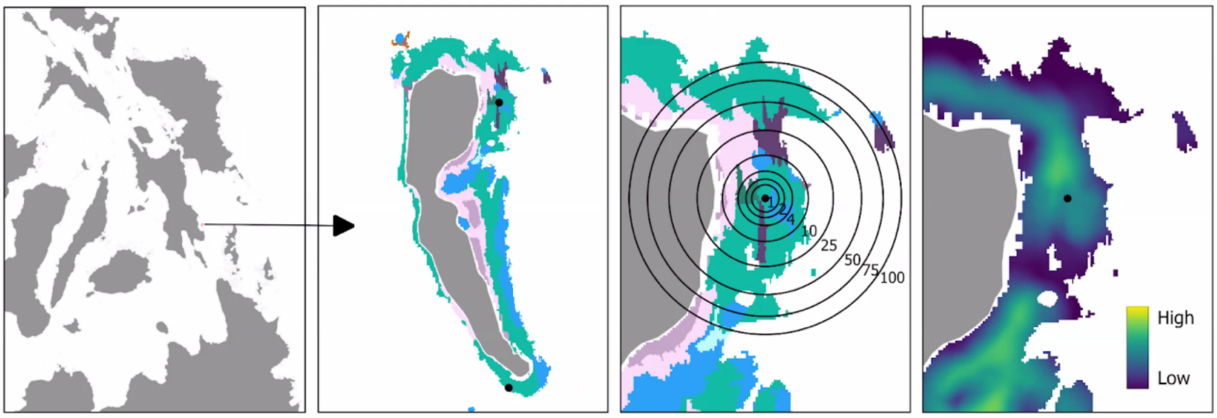


**Figure S6** A schematic representation of the steps involved in estimating habitat heterogeneity. First, a map of benthic habitat types or geomorphic zones is chosen for a given geographic region. Then, the diversity of benthic habitat or geomorphic classes is calculated using Shannon's diversity index or Rao's Q within defined spatial windows. The resulting output provides a means of summarising habitat heterogeneity at various spatial scales at each sampling location.

*1.4.3 Temperature metrics*

Daily values were obtained for each site by averaging over 24 hours starting at midnight. Daily temperature range was calculated as the difference between the highest and lowest temperature each day. Mean daily estimates were obtained by averaging over the 2021-2023 period. Daily temperatures were further averaged across months to obtain mean monthly estimates. All daily and monthly values were averaged per site to obtain mean metrics.

**Table S2** An overview of the various temperature measures and metrics calculated and used in this study from *in situ* logger data.

| *Metric* | *Time scale* | *Relevance* | *Used in DAG* |
|---|---|---|---|
| Daily mean temperature | Daily | Daily averages | No |
| Daily temperature range | Daily | Daily fluctuations | No |
| Mean monthly temperatures | Monthly | Monthly patterns | No |
| Maximum monthly temperatures | Monthly | Thermal extremes | No |
| Minimum monthly temperatures | Monthly | Thermal extremes | No |
| Mean daily temperature | Entire period | Average thermal conditions | Yes: mean temperature |
| Standard deviation in monthly temperatures | Entire period | Seasonal variation | Yes: temperature variation |
| Thermal predictability | Entire period | Pattern regularity | Yes: thermal predictability |

*1.4.4 Autocorrelation amongst variables*

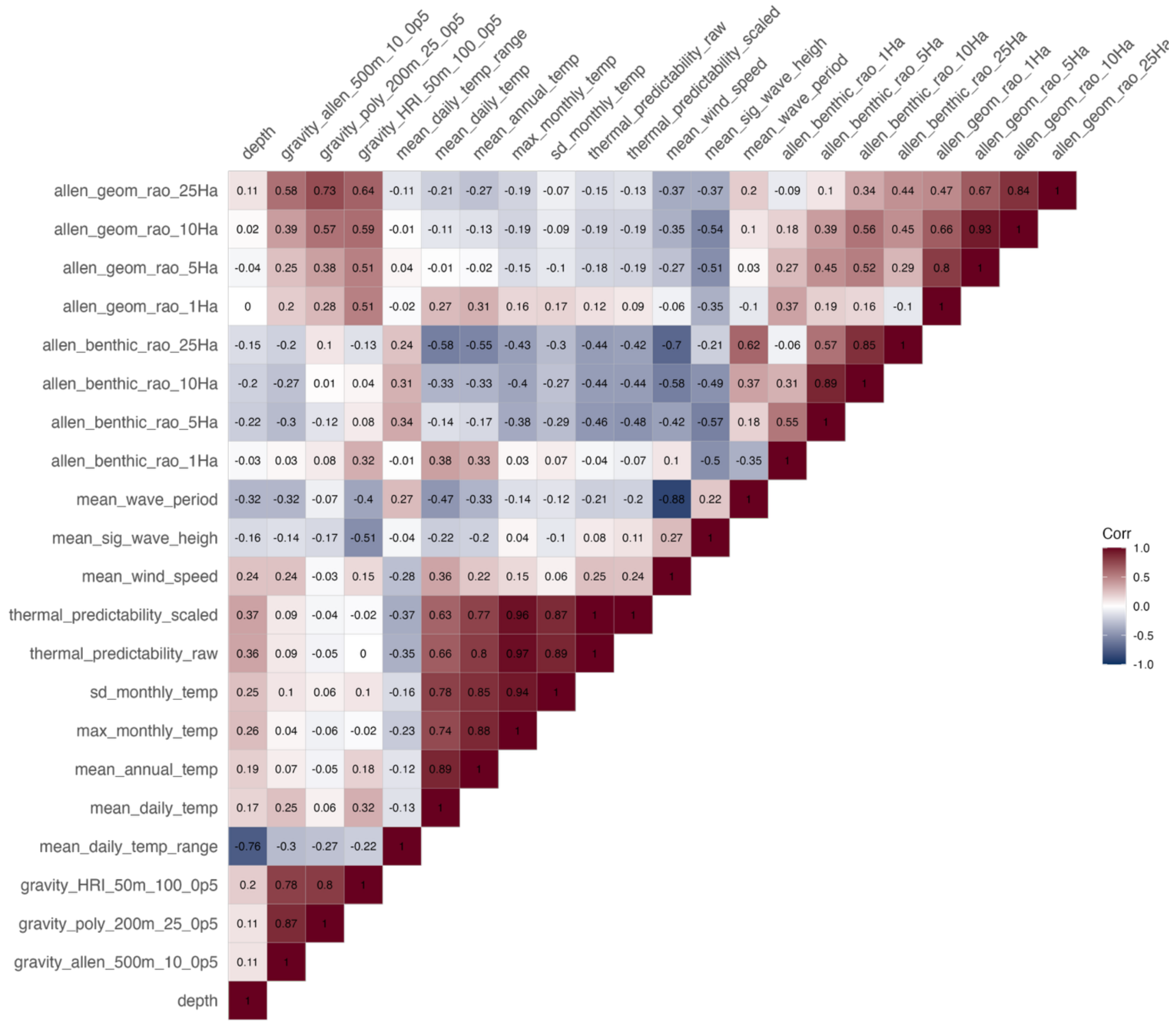


**Figure S7** Pearson's pairwise correlation matrix of environmental covariates. Environmental predictors with Pearson's coefficients > 0.6 were never included in the same model.

1.5 Structural Equation Modelling

We specified a directed acyclic graph (DAG) representing hypothesized causal relationships among environmental variables and genetic diversity (Fig. 1 in the main text). This theoretical structure was based on *a priori* theoretical expectations and empirical evidence and was used to identify minimal adjustment sets for causal inference. However, minimal adjustment sets are influenced by conditional dependencies or independencies among variables, which can be validated using empirical data.

Structural equation modelling (SEM) was conducted using the R package piecewiseSEM v2.3.0.1 to validate expected conditional independence amongst variables. Separate structural equation models were run for the aggregate and species-specific datasets. Relationships were tested using simple linear models and as such random effects or hierarchical data (e.g., multiple individuals per sampling location) could not be incorporated so models were run at the population/location level. Population observed heterozygosity was used as the response variable, and seascape variables included in the model as predictors comprised: reef gravity, habitat heterogeneity (benthic and geomorphic), temperature (mean daily and monthly standard deviation), thermal predictability, and wave energy (mean wave period and mean significant wave height). Where multiple proxies were available for a single exposure variable, i.e. temperature, models were tested with different combinations of each focal seascape variable. Geographic distance was incorporated in models by converting the geographic distance matrix to Moran Eigenvector Maps (MEMs) and including the first MEM (up to three MEMs were included in exploratory model iterations). While latent variables unmeasured variables were kept in the DAG, their effect on other variables could not be assessed using SEM and as such we only tested a reduced model. In this reduced model, we expected wave exposure and other environmental variables to show ‘direct’ effects on genetic diversity that may represent pathways mediated by unmeasured variables such as population size and structural complexity.

**Results**

2.1 Caribbean *Agaricia spp.* delineation

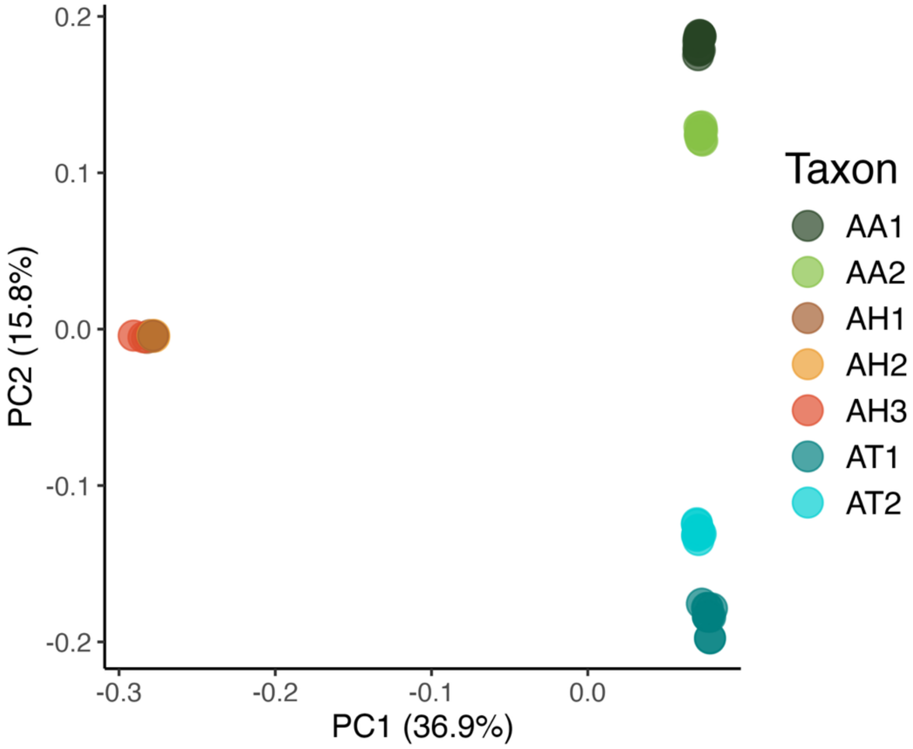


**Figure S8** Principal Components Analysis (PCA) showing the clear genetic differentiation of both *Agaricia tenuifolia* lineages identified in this study from other *Agaricia spp.* found in the Caribbean. Data corresponding to AA and AH were generated by Prata et al. (2024), with samples collected in Curaçao. Codes: AA – *Agaricia agaricites*, AH – *Agaricia humilis*, AT – *Agaricia tenuifolia*.

### 2.2 *Agaricia tenuifolia* cryptic lineages

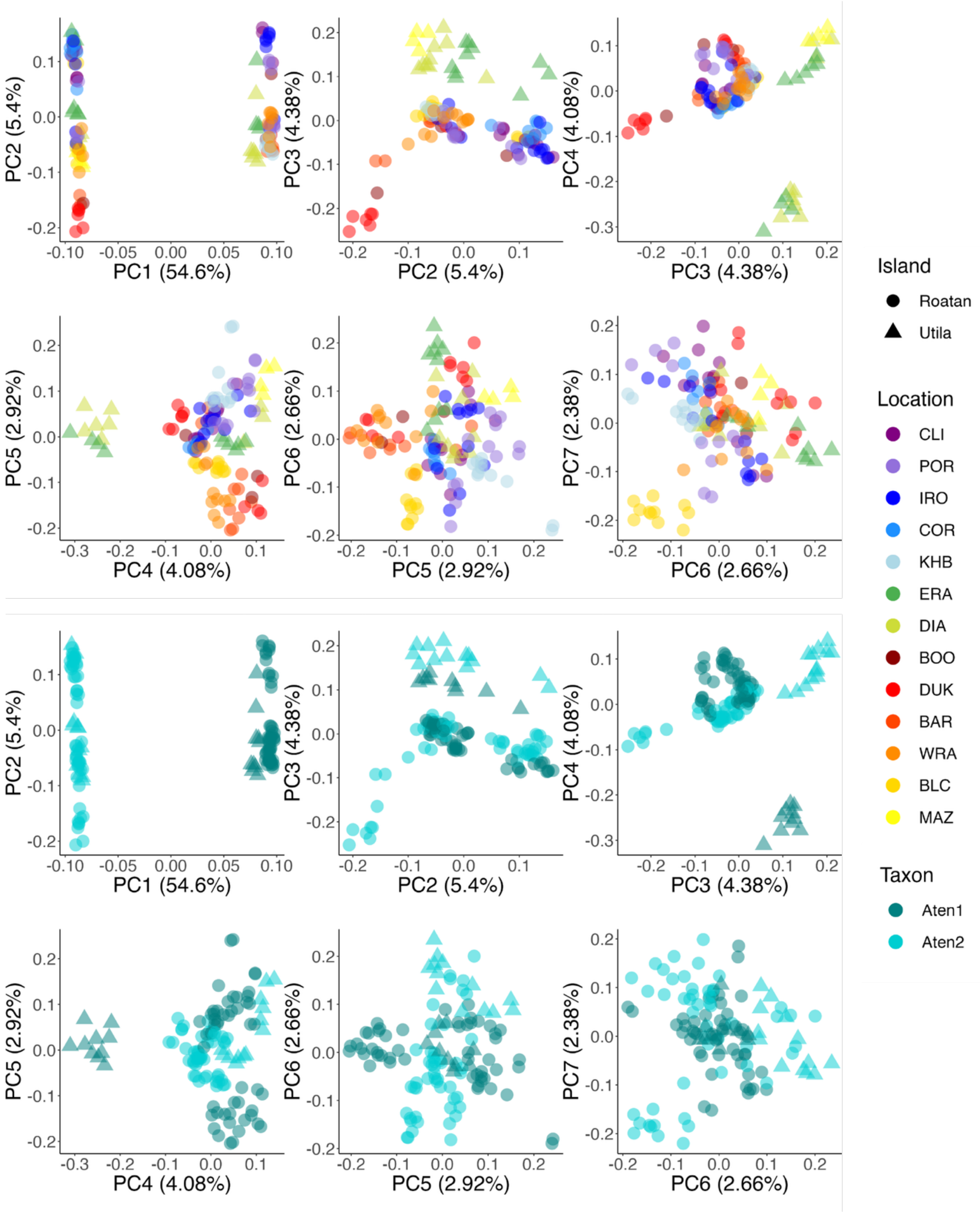


**Figure S9** Principal Components axes one and two clearly show the genetic separation between AT1 and AT2 (54% of the total genetic variation is captured by PC1), while other PC axes (two – five) emphasise the geographic structure within genotypes. Principal Components Analysis (PCA) plots were generated using ddRAD genotype information, with PC axes one to six shown. Individuals are coloured by location (top) and cryptic species assignment (bottom).

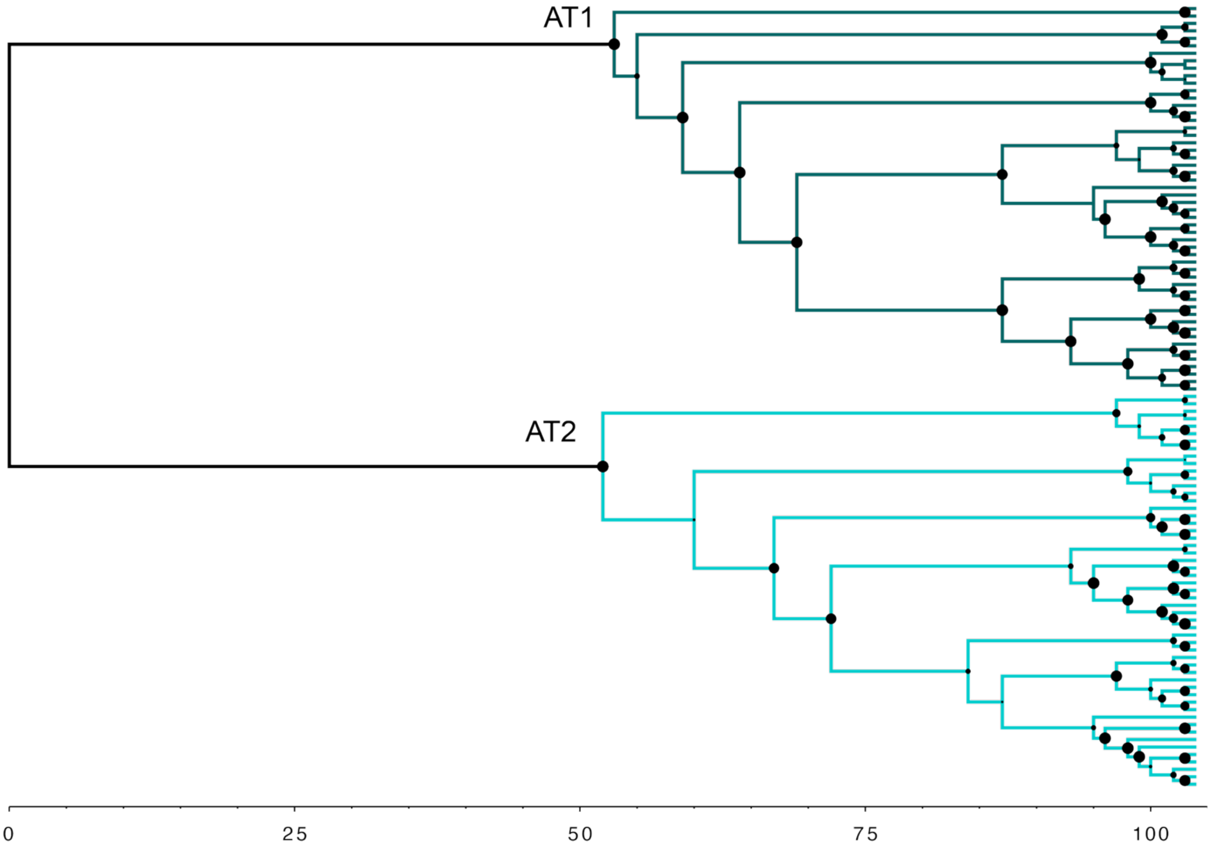


**Figure S10** Phylogenetic tree generated using ddRAD SNP data analysed using IQtree and visualised using FigTree v1.4, supports the presence of two delineated clades. Size of circle at each branch node denotes the magnitude of bootstrap support (larger circles are closer to 100% support, with values ranging from 43–100%).

2.3 Intraspecific analyses

**Table S3** Genome-wide diversity metrics generated using STACKS for the two *Agaricia spp.* assessed in this study. Sample numbers per population were varied to assess resultant changes in observed heterozygosity and nucleotide diversity estimates. Estimates for all populations within a given species were averaged to produce the summary values listed below.

| *Taxon* | *Number of individuals per population* | *Mean nucleotide diversity* | *Mean nucleotide diversity (variant)* | *Mean observed heterozygosity* | *Mean observed heterozygosity (variant)* |
|---|---|---|---|---|---|
| AT1 | 2 | 0.0033 | 0.201 | 0.0024 | 0.151 |
| | 3 | 0.0036 | 0.228 | 0.0024 | 0.156 |
| | 5 | 0.0035 | 0.234 | 0.0023 | 0.161 |
| AT2 | 2 | 0.0038 | 0.176 | 0.0024 | 0.114 |
| | 3 | 0.0037 | 0.185 | 0.0024 | 0.129 |
| | 5 | 0.0043 | 0.264 | 0.0025 | 0.158 |

**Table S4** Comparison of genome-wide diversity metrics generated using STACKS (n = 2 per population) and custom scripts (individual-level estimates). Estimates were only generated if there were enough individuals per population/location. Individual-level estimates were only calculated for observed heterozygosity and were averaged per sampling location to produce the summary values below (Mean $H_o$). Genetic diversity estimates were generated using both invariant and variant sites, or only variant sites. Empty rows correspond to populations for which there were not enough individuals. Pop: population or sampling location, *Ho*: observed heterozygosity, *Pi*: nucleotide diversity, SNP: estimates calculated using variant sites only.

| | | **n = 3** | | | | **n = 1** | |
|---|---|---|---|---|---|---|---|
| *Taxon* | *Pop* | $H_o$ | $H_o$ *(SNP)* | *Pi* | *Pi (SNP)* | *Mean* $H_o$ | *Mean* $H_o$ *(SNP)* |
| AT1 | BAR1 | 0.00238 | 0.1368 | 0.00459 | 0.26415 | 0.001535 | 0.07351 |
| | BAR2 | | | | | 0.001874 | 0.084619 |
| | BOO1 | 0.00271 | 0.17467 | 0.00364 | 0.23416 | 0.001801 | 0.080901 |
| | DIA1 | 0.00257 | 0.14824 | 0.00473 | 0.27309 | 0.001329 | 0.076314 |
| | DUK2 | | | | | 0.001514 | 0.070244 |
| | ERA1 | | | | | 0.002743 | 0.141276 |
| | IRO1 | | | | | 0.001923 | 0.114637 |
| | IRO2 | 0.0026 | 0.16575 | 0.00339 | 0.2159 | 0.001633 | 0.07034 |
| | KHB1 | 0.00222 | 0.13018 | 0.00256 | 0.15031 | 0.001657 | 0.066772 |
| | KHB2 | 0.00243 | 0.16399 | 0.00332 | 0.22419 | 0.001682 | 0.064817 |
| | POR1 | 0.00239 | 0.14368 | 0.00269 | 0.16211 | 0.00183 | 0.069444 |
| | POR2 | 0.00222 | 0.15146 | 0.00248 | 0.16911 | 0.001691 | 0.066087 |

| | | | | | | | |
|---|---|---|---|---|---|---|---|
| | WRA1 | 0.00233 | 0.14217 | 0.00259 | 0.15816 | 0.001611 | 0.063521 |
| | WRA2 | 0.00242 | 0.15064 | 0.00256 | 0.15926 | 0.001622 | 0.064828 |
| AT2 | BLC1 | 0.00221 | 0.10681 | 0.00372 | 0.17987 | 0.001484 | 0.040039 |
| | BLC2 | 0.00248 | 0.13464 | 0.00357 | 0.1939 | 0.001568 | 0.041179 |
| | BOO1 | | | | | 0.001264 | 0.034337 |
| | CLI1 | 0.00243 | 0.08548 | 0.00362 | 0.12715 | 0.001366 | 0.0386 |
| | CLI2 | 0.00223 | 0.11278 | 0.00392 | 0.19787 | 0.001447 | 0.039244 |
| | COR1 | 0.00291 | 0.12144 | 0.00357 | 0.14934 | 0.001633 | 0.043452 |
| | COR2 | 0.00299 | 0.11286 | 0.00543 | 0.20508 | 0.001574 | 0.042896 |
| | DIA2 | 0.00231 | 0.10825 | 0.00359 | 0.16786 | 0.001218 | 0.03455 |
| | DUK1 | 0.00219 | 0.12005 | 0.00383 | 0.20973 | 0.001866 | 0.052661 |
| | DUK2 | 0.00228 | 0.10653 | 0.00379 | 0.17669 | 0.001404 | 0.041043 |
| | ERA1 | 0.0023 | 0.0789 | 0.00369 | 0.12647 | 0.00103 | 0.029539 |
| | ERA2 | 0.00283 | 0.12323 | 0.00873 | 0.38053 | 0.001594 | 0.043664 |
| | IRO1 | 0.00233 | 0.09443 | 0.00383 | 0.15509 | 0.001273 | 0.034546 |
| | IRO2 | 0.00247 | 0.12983 | 0.00298 | 0.15669 | 0.000846 | 0.020786 |
| | MAZ2 | 0.00231 | 0.14451 | 0.00251 | 0.15666 | 0.001094 | 0.031421 |
| | POR1 | | | | | 0.000972 | 0.024988 |
| | POR2 | 0.00213 | 0.11553 | 0.00242 | 0.13146 | 0.001043 | 0.026462 |
| | WRA1 | 0.0023 | 0.10297 | 0.00247 | 0.11069 | 0.000855 | 0.026869 |
| | WRA2 | 0.00224 | 0.13851 | 0.00258 | 0.1596 | 0.000836 | 0.026661 |

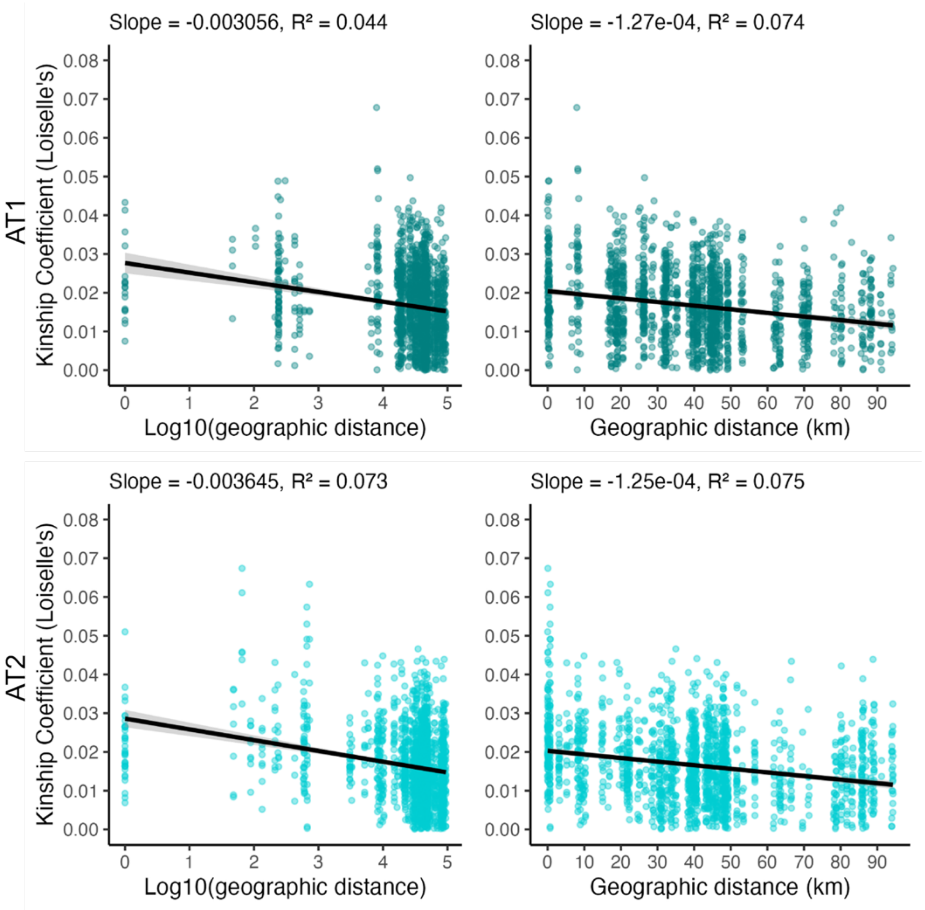


**Figure S11** Isolation-by-Distance (IBD) slopes generated by plotting Loiselle's *F* as a measure of genetic distance (estimated using SPAGeDi) against geographic distance.

**Table S5** Estimates of neighbourhood size (*NS*) and dispersal variance (*σ,* sigma). Genepop was used to estimate *NS* and *σ* based on Rousset's *â*. Effective population density relates to the number of individuals per square distance unit. SPAGeDi was used to estimate *NS* and *σ* based on Loiselles's *F*, assuming a 2-dimensional (2D) population at drift-dispersal equilibrium under isotropic dispersal. SPAGeDi outputs standard errors for estimates by jack-knifing over loci. Estimation based on the regression slope between sigma and 20 sigma following an iterative procedure. *NS* for AT1 is 579.70 (SE = 22.49), and 628.80 (SE = 21.38) for AT2 under a 2D model based on Rousset's *â*. For Loiselle's *F*, NS estimates varied with De and are shown below. Analyses included 61 individuals for AT1 and 63 individuals for AT2. Refer to Figure S10 for examples of the IbD regression slopes generated using Loiselle's *F*. † SPAGeDi iterations for which convergence was not achieved when removing some loci during jack-knifing procedure.

| | ***Effective population density*** | ***Neighbourhood and σ based on Rousset's â*** | | ***Neighbourhood and σ based on Loiselle's F*** | | | |
|---|---|---|---|---|---|---|---|
| *Taxon* | *$D_e$ ($m^2$)* | *$σ_e$ (m)* | *SE* | *NS* | *SE* | *$σ_e$ (m)* | *SE* |
| AT1 | 0.02 | 13.55 | 0.2628 | 231.192 | 93.4161 | 30.3301 | 6.08092 |
| | 0.04 | 9.58 | 0.1858 | 222.693 | 364.46 | 21.0487 | 16.3945 |
| | 0.06 | 7.82 | 0.1517 | 171.981 | 658.613 | 15.1031 | 26.784 |
| | 0.08 | 6.77 | 0.1314 | 171.981 | 241.882 | 13.0797 | 9.46303 |
| | 0.1 | 6.06 | 0.1175 | † | † | † | † |
| | 0.12 | 5.53 | 0.1073 | † | † | † | † |
| | 0.14 | 5.12 | 0.0993 | † | † | † | † |
| | 0.16 | 4.79 | 0.0929 | † | † | † | † |
| | 0.18 | 4.52 | 0.0876 | † | † | † | † |
| | 0.2 | 4.28 | 0.0831 | † | † | † | † |
| | 0.3 | 3.50 | 0.0678 | † | † | † | † |
| | 0.4 | 3.03 | 0.0588 | † | † | † | † |
| AT2 | 0.02 | 14.11 | 0.2399 | † | † | † | † |
| | 0.04 | 9.98 | 0.1696 | † | † | † | † |
| | 0.06 | 8.15 | 0.1385 | 192.054 | 90.507 | 15.9602 | 3.75058 |
| | 0.08 | 7.06 | 0.1199 | 192.054 | 90.507 | 13.8219 | 3.24809 |
| | 0.1 | 6.31 | 0.1073 | 192.054 | 89.946 | 12.3627 | 2.88624 |
| | 0.12 | 5.76 | 0.0979 | 179.751 | 136.889 | 10.9181 | 4.6281 |
| | 0.14 | 5.33 | 0.0907 | 79.8705 | 130.05 | 6.73799 | 4.86057 |
| | 0.16 | 4.99 | 0.0848 | 104.444 | 62.657 | 7.15671 | 2.0179 |
| | 0.18 | 4.70 | 0.0800 | 104.444 | 62.657 | 6.74741 | 1.9025 |
| | 0.2 | 4.46 | 0.0758 | 104.444 | 62.657 | 6.40115 | 1.80487 |
| | 0.3 | 3.64 | 0.0619 | 129.017 | 108.4 | 5.85012 | 2.44873 |
| | 0.4 | 3.16 | 0.0536 | 129.017 | 108.4 | 5.06635 | 2.12065 |

**Table S6** Comparison of 1-dimensional (1D) and 2-dimensional (2D) geographic model outputs using genepop. *NS* for AT1 is 46.13 (SE = 1.79) under the 1D model, and 579.70 (SE = 22.49) under the 2D model. *NS* for AT2 is 50.04 (SE = 1.70) under the 1D model, and 628.80 (SE = 21.38) under the 2D model. Analyses included 61 individuals for AT1 and 63 individuals for AT2.

| *Taxon* | *$D_e$ ($m^2$)* | *σ* | *σ SE* | *σ* | *σ SE* |
|---|---|---|---|---|---|
| AT1 | 0.02 | 33.96 | 0.6587 | 13.55 | 0.2628 |
| | 0.04 | 24.01 | 0.4658 | 9.58 | 0.1858 |
| | 0.06 | 19.61 | 0.3803 | 7.82 | 0.1517 |
| | 0.08 | 16.98 | 0.3293 | 6.77 | 0.1314 |
| | 0.1 | 15.19 | 0.2946 | 6.06 | 0.1175 |
| | 0.12 | 13.86 | 0.2689 | 5.53 | 0.1073 |
| | 0.14 | 12.84 | 0.2490 | 5.12 | 0.0993 |
| | 0.16 | 12.01 | 0.2329 | 4.79 | 0.0929 |
| | 0.18 | 11.32 | 0.2196 | 4.52 | 0.0876 |
| | 0.2 | 10.74 | 0.2083 | 4.28 | 0.0831 |
| | 0.3 | 8.77 | 0.1701 | 3.50 | 0.0678 |
| | 0.4 | 7.59 | 0.1473 | 3.03 | 0.0588 |
| | 0.6 | 6.20 | 0.1203 | 2.47 | 0.0480 |
| | 0.8 | 5.37 | 0.1041 | 2.14 | 0.0415 |
| | 1.0 | 4.80 | 0.0932 | 1.92 | 0.0372 |
| AT2 | 0.02 | 35.37 | 0.6012 | 14.11 | 0.2399 |
| | 0.04 | 25.01 | 0.4251 | 9.98 | 0.1696 |
| | 0.06 | 20.42 | 0.3471 | 8.15 | 0.1385 |
| | 0.08 | 17.68 | 0.3006 | 7.06 | 0.1199 |
| | 0.1 | 15.82 | 0.2689 | 6.31 | 0.1073 |
| | 0.12 | 14.44 | 0.2454 | 5.76 | 0.0979 |
| | 0.14 | 13.37 | 0.2272 | 5.33 | 0.0907 |
| | 0.16 | 12.50 | 0.2126 | 4.99 | 0.0848 |
| | 0.18 | 11.79 | 0.2004 | 4.70 | 0.0800 |
| | 0.2 | 11.18 | 0.1901 | 4.46 | 0.0758 |
| | 0.3 | 9.13 | 0.1552 | 3.64 | 0.0619 |
| | 0.4 | 7.91 | 0.1344 | 3.16 | 0.0536 |
| | 0.6 | 6.46 | 0.1098 | 2.58 | 0.0438 |
| | 0.8 | 5.59 | 0.0951 | 2.23 | 0.0379 |
| | 1.0 | 5.00 | 0.0850 | 2.00 | 0.0339 |

### 2.4 Environmental variable optimisation

#### *2.4.1 Reef gravity optimisation*

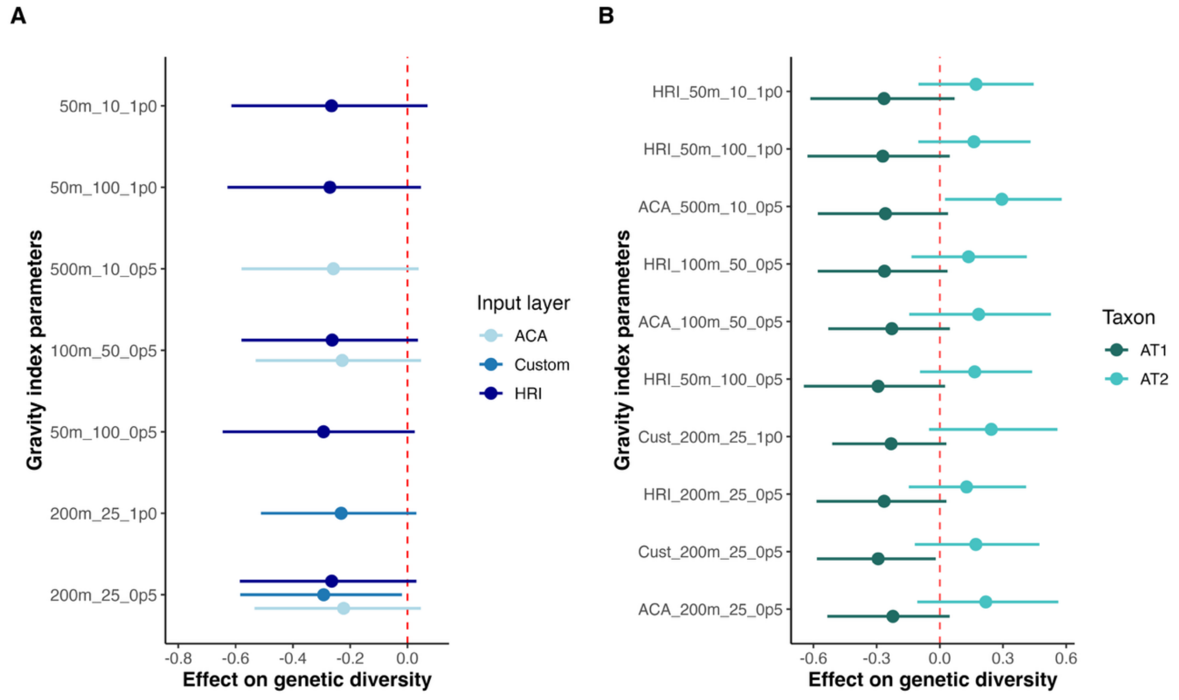


**Figure S12** Posterior mean values and distributions (standardised) associated with the effect of various gravity metrics on genetic diversity revealed the '200m_25_0p5' resolution metric calculated using the custom reef polygons as the best gravity metric.

#### *2.4.2 Habitat heterogeneity optimisation*

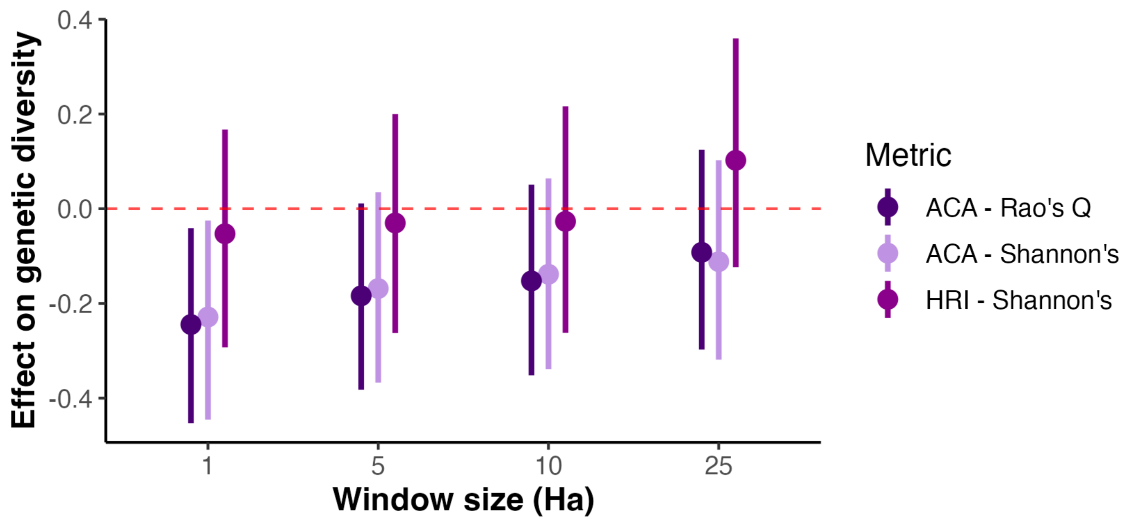


**Figure S13** Posterior mean values and distributions (standardised) associated with the effect of benthic habitat heterogeneity on genetic diversity showed the strongest effects at 1 Ha window sizes using the Allen Coral Atlas map, regardless of the mathematical equation (Rao's Q versus Shannon's diversity) used.

2.5 Causal framework

*2.5.1 Minimum adjustment sets*

The minimal adjustment sets resulting from our Directed Acyclic Graph (DAG) informed our model syntax and covariate inclusion for testing the effect of each exposure variable on genetic diversity. Notably, testing the effect of thermal predictability on genetic diversity required the inclusion of a latent variable (structural complexity) and as such thermal predictability was not reported as a focal variable in the main text. However, exploratory analyses assessing the effect of thermal predictability on genetic diversity are reported below in sections 2.5.2 and 2.9.

**Table S7** Minimum adjustment sets based on the Directed Acyclic Graph described in this study. Adjustment sets represent the specific variables that should be accounted for in models assessing the relationship between the designated exposure variable and genetic diversity. Where no adjustment set is specified, only the exposure variable needs to be included in statistical models. The theoretical DAG corresponds to the one shown in the main text. The proxy DAG corresponds to one where reef gravity is included as a proxy for reef area and connectivity.

| *DAG version* | *Exposure variable* | *Adjustment set* |
|---|---|---|
| Theoretical | Reef gravity | NA |
| | Habitat heterogeneity | Wave exposure |
| | Mean temperature | Wave exposure |
| | Thermal predictability/temperature variation | Wave exposure + habitat heterogeneity + structural complexity |
| Proxy | Reef gravity | none |
| | Habitat heterogeneity | Wave exposure |
| | Mean temperature | Wave exposure |
| | Temperature predictability/temperature variation | Wave exposure + habitat heterogeneity + structural complexity |

Conditional independence checks revealed that thermal predictability and mean temperature shared a causal pathway. However, this is unsurprising given that both metrics were highly correlated based on a Pearson's correlation coefficient (Fig. S7). Redrawing the DAG with the inclusion of a casual pathway connecting these temperature metrics did not affect the minimal adjustment sets outlined above. Moreover, for the aggregate and AT2 datasets, wave exposure (mean wave period) significantly influenced genetic diversity (pop-$H_o$). We updated our DAG based on the SEM findings that thermal predictability significantly influenced mean temperature ($\beta = 0.63 - 0.67$, $p < 0.001$ across models) and mean wave period influenced genetic diversity ($\beta = -0.54 - -0.68$, $p < 0.001$). This revision required adjusting for thermal predictability when estimating the effect of mean temperature on genetic diversity but did not influence any other minimal adjustment sets. However, because thermal predictability and mean temperature were highly correlated ($r = 0.63$, Fig. S7) they were not included together in the Bayesian models reported in the main text. We did however run structural equation models with both variables included as well as conduct separate tests for each temperature metric to avoid multicollinearity (see next section).

*2.5.2 Structural Equation Modelling*

Structural equation modelling (SEM) revealed largely congruent results to our Bayesian GLMMs indicating that our main conclusions are robust to different modelling approaches (Fig. S13). SEM results after accounting for non-independence amongst variables revealed our causal structure was consistent with the empirical data despite some hypothesized pathways showing weak or non-significant effects (likely due to limited sample sizes of 14-24 populations).

Consistent with our GLMM results, reef gravity was significantly associated with lower genetic diversity in AT1. Benthic heterogeneity (1 Ha window size) showed negative relationships with genetic diversity in both species; however, these effects were marginal ($p = 0.092$ and $p = 0.088$ respectively), likely reflecting reduced statistical power at the population scale. The consistent direction and magnitude of effects across analytical approaches suggests benthic heterogeneity may influence genetic diversity, though additional sampling would strengthen this inference. Moreover, mean temperature was associated with lower genetic diversity across all models, with the strongest effect seen for AT2 (although the effect was significant in the SEM but marginal in the GLMM). Thermal predictability was mostly positively correlated with genetic diversity, but all associations were non-significant.

Interestingly, for the aggregate and AT2 datasets, wave period showed a strong direct effect on genetic diversity, while other environmental variables showed weaker or non-significant direct effects. We interpret this as consistent with our theoretical model wherein wave exposure influences genetic diversity through effects on population size and demography, which could not be directly measured in this study. Future studies incorporating population census data would help partition direct versus population-mediated effects of wave exposure on genetic diversity.

**Table S8** Coefficients and p-values (***p < 0.001, **p < 0.01, *p < 0.05) resulting from SEMs conducted with the entire dataset (n = 24, $R^2 = 0.41$). Variables are listed in order of importance i.e., most significant effect on the response.

| *Predictor* | *Direct Effect (β)* | *P-value* | *Indirect Effects* | *Total Effect* |
|---|---|---|---|---|
| **Wave period** | -0.540 | 0.020 * | -0.096 | -0.636 |
| via benthic het | - | - | -0.353 × -0.297 = 0.105 | - |
| via thermal pred | - | - | -0.260 × 0.463 = -0.120 | - |
| via mean temp | - | - | -0.224 × -0.413 = 0.093 | - |
| via benthic het → temp | - | - | -0.353 × 0.328 × -0.413 = 0.048 | - |
| via thermal pred → temp | - | - | -0.260 × 0.626 × -0.413 = 0.067 | - |
| **Benthic heterogeneity** | -0.297 | 0.192 | -0.084 | -0.381 |
| via thermal pred | - | - | -0.130 × 0.463 = -0.060 | - |
| via mean temp | - | - | 0.328 × -0.413 = -0.135 | - |
| via thermal pred → temp | - | - | -0.130 × 0.626 × -0.413 = 0.034 | - |
| **Thermal predictability** | 0.463 | 0.102 | -0.259 | 0.204 |
| via mean temp | - | - | 0.626 × -0.413 = -0.259 | - |
| **Mean temperature** | -0.413 | 0.194 | 0 | -0.413 |
| **Reef gravity** | -0.074 | 0.692 | 0 | -0.074 |

**Table S9** Coefficients and p-values (***p < 0.001, **p < 0.01, *p < 0.05) resulting from SEMs conducted with the AT1 dataset (n = 14, $R^2 = 0.86$). Spatial MEMs were included in this model i.e., MEM1 is the first Moran's Eigenvector Map capturing dominant spatial autocorrelation patterns. Variables are listed in order of importance.

| *Predictor* | *Direct Effect (β)* | *P-value* | *Indirect Effects* | *Total Effect* |
|---|---|---|---|---|
| **Habitat connectivity** | -0.947 | 0.005 * | -0.010 | -0.957 |
| via thermal pred | - | - | 0.140 × -0.139 = -0.019 | - |
| via mean temp | - | - | 0.109 × -0.155 = -0.017 | - |
| via thermal pred → temp | - | - | 0.140 × -0.013 × -0.155 = 0.000 | - |
| **Benthic heterogeneity** | -0.614 | 0.092 | 0.078 | -0.536 |
| via thermal pred | - | - | 0.139 × -0.139 = -0.019 | - |

| | | | | |
|---|---|---|---|---|
| via mean temp | - | - | 0.714 × -0.155 = -0.111 | - |
| via thermal pred → temp | - | - | 0.139 × -0.013 × -0.155 = 0.000 | - |
| **Wave period** | 0.037 | 0.891 | 0.001 | 0.038 |
| via benthic het | - | - | -0.152 × -0.614 = 0.093 | - |
| via thermal pred | - | - | -0.413 × -0.139 = 0.057 | - |
| via mean temp | - | - | -0.025 × -0.155 = 0.004 | - |
| all indirect pathways | - | - | -0.092 | - |
| **Thermal predictability** | -0.139 | 0.733 | 0.002 | -0.137 |
| Via mean temp | - | - | -0.013 × -0.155 = 0.002 | - |
| **Mean temperature** | -0.155 | 0.536 | 0 | -0.155 |
| **MEM1** | 0.887 | 0.102 | -0.058 | 0.829 |
| via benthic het | - | - | 0.620 × -0.614 = -0.381 | - |
| via thermal pred | - | - | -0.999 × -0.139 = 0.139 | - |
| via mean temp | - | - | Not significant | - |
| **MEM2** | -0.357 | 0.300 | 0.007 | -0.350 |
| via thermal pred | - | - | 0.544 × -0.139 = -0.076 | - |

**Table S10** Coefficients and p-values (***p < 0.001, **p < 0.01, *p < 0.05) resulting from SEMs conducted with the AT2 dataset (n = 18, $R^2$ = 0.67). Variables are listed in order of importance.

| *Predictor* | *Direct Effect (β)* | *P-value* | *Indirect Effects* | *Total Effect* |
|---|---|---|---|---|
| **Wave period** | -0.678 | 0.003 * | -0.155 | -0.833 |
| via benthic het | - | - | -0.097 × -0.324 = 0.031 | - |
| via thermal pred | - | - | -0.192 × 0.438 = -0.084 | - |
| via mean temp | - | - | -0.401 × -0.761 = 0.305 | - |
| via benthic het → temp | - | - | -0.097 × 0.206 × -0.761 = 0.015 | - |
| **Mean temperature** | -0.761 | 0.014 * | 0 | -0.761 |
| **Benthic heterogeneity** | -0.324 | 0.088 | -0.157 | -0.481 |
| via thermal pred | - | - | -0.100 × 0.438 = -0.044 | - |
| via mean temp | - | - | 0.206 × -0.761 = -0.157 | - |
| **Thermal predictability** | 0.438 | 0.095 | 0 | 0.438 |
| **Reef gravity** | 0.075 | 0.661 | 0 | 0.075 |

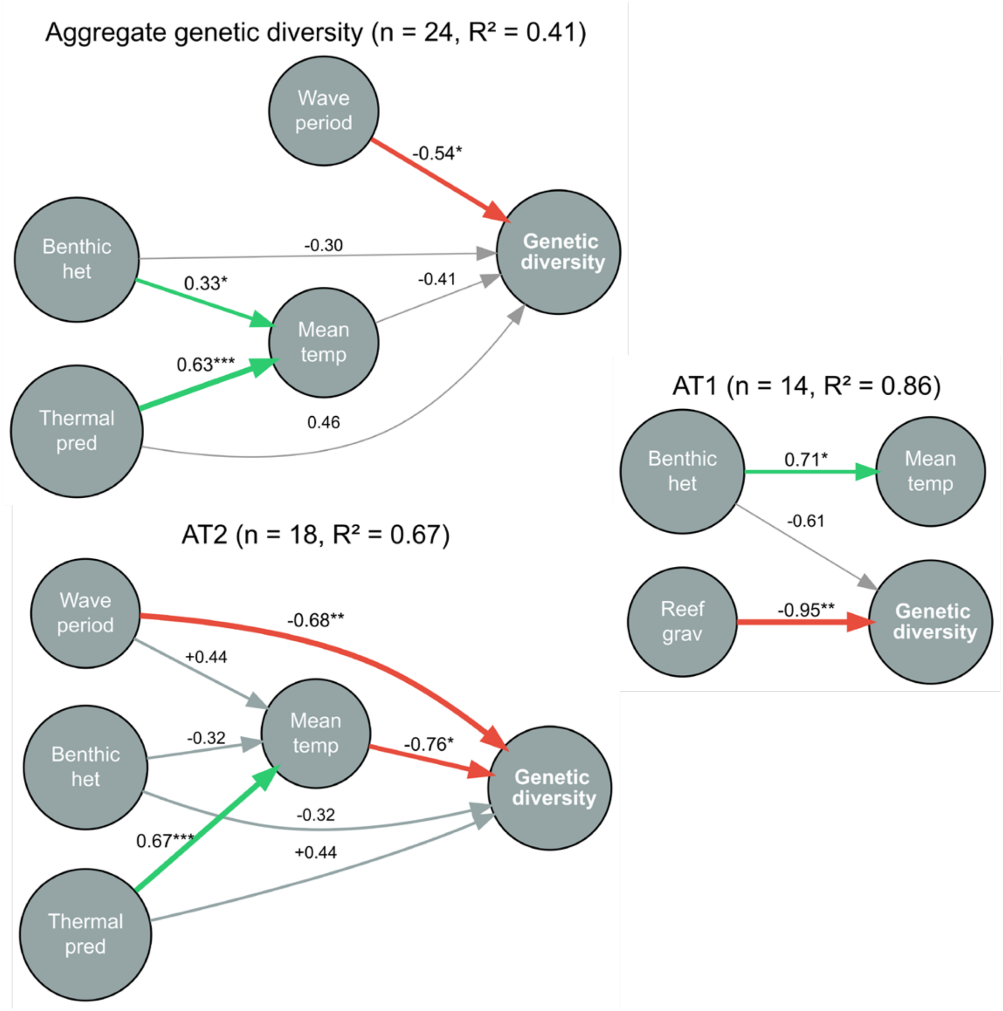


**Figure S14** Structural Equation Modelling (SEM) results examining relationships between environmental predictors and genetic diversity in two coral species. $R^2$ values represent the proportion of variance explained in endogenous variables, while n represents the number of populations included in the model. Standardised path coefficients (β) are shown above each arrow. Thick arrows denote significant pathways based on a p-value threshold of 0.05 (***p < 0.001, **p < 0.01, *p < 0.05), while grey arrows symbolise marginal pathways (0.05 < p < 0.1). Red and green arrows indicate negative and positive effects, respectively.

### 2.6 Model comparisons

#### *2.6.1 Individual versus population response*

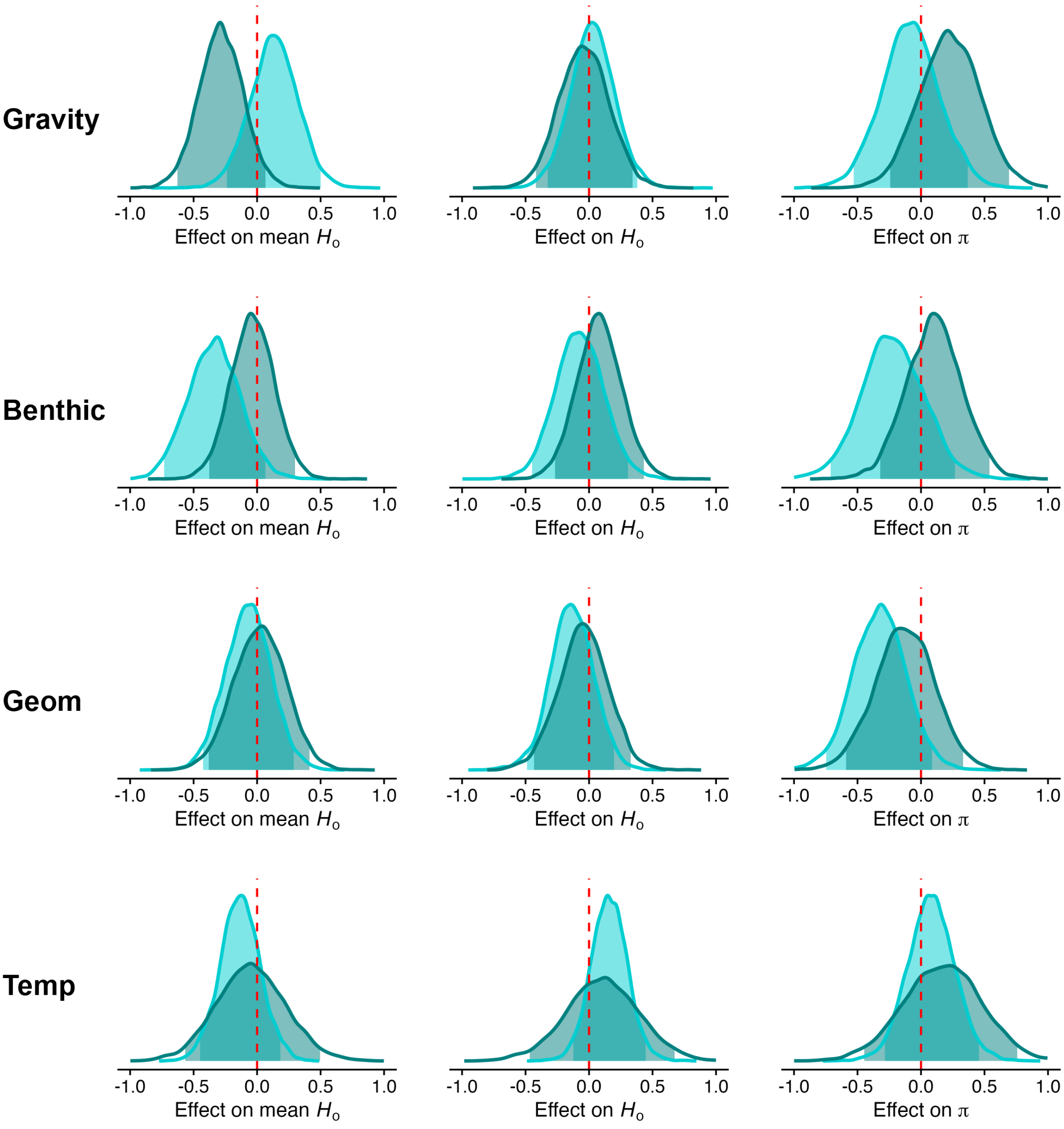


**Figure S15** Comparison of brms models assessing the relationship between reef gravity (Gravity), benthic (1 Ha) heterogeneity (Benthic), geomorphic (1 Ha) heterogeneity (Geom) and temperature (Temp) and three estimates of population-level genetic diversity reveal similar patterns, but disparate effect sizes. Mean $H_o$: mean individual-level genetic diversity i.e., average of individual-level values used in models presented in the main text, $H_o$: observed heterozygosity generated using STACKS, $\pi$: nucleotide diversity (pi) generated using STACKS. All measures of genetic diversity were estimated considering both variant and invariant sites. Individuals were assigned to populations based on sampling location e.g., BAR1 and ERA2.

*2.6.2 Effect of temperature on genetic diversity*

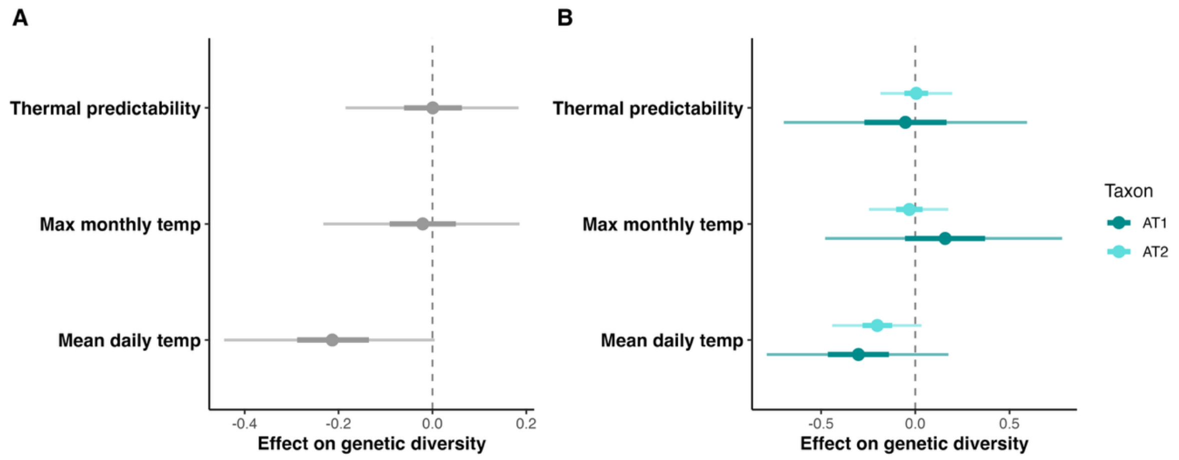


**Figure S16** Comparison of the effect of different temperature metrics on genetic diversity reveals the strongest effect when mean daily temperature is used as the focal predictor of both overall and species-specific genetic diversity (individual $H_o$). A) Overall model that does not allow slopes to differ between species. B) Species-specific model that allows for differences in posterior mean distributions between species.

*2.6.3 Spatially explicit models*

Congruent with the models reported in the main text, INLA models (Fig. S15) revealed significant associations between reef gravity and genetic diversity in AT1, and benthic heterogeneity and genetic diversity in AT2. Moreover, geomorphic was found to be significantly associated with lower nucleotide diversity in AT2. All other relationships were associated with wide credible intervals that overlapped with zero.

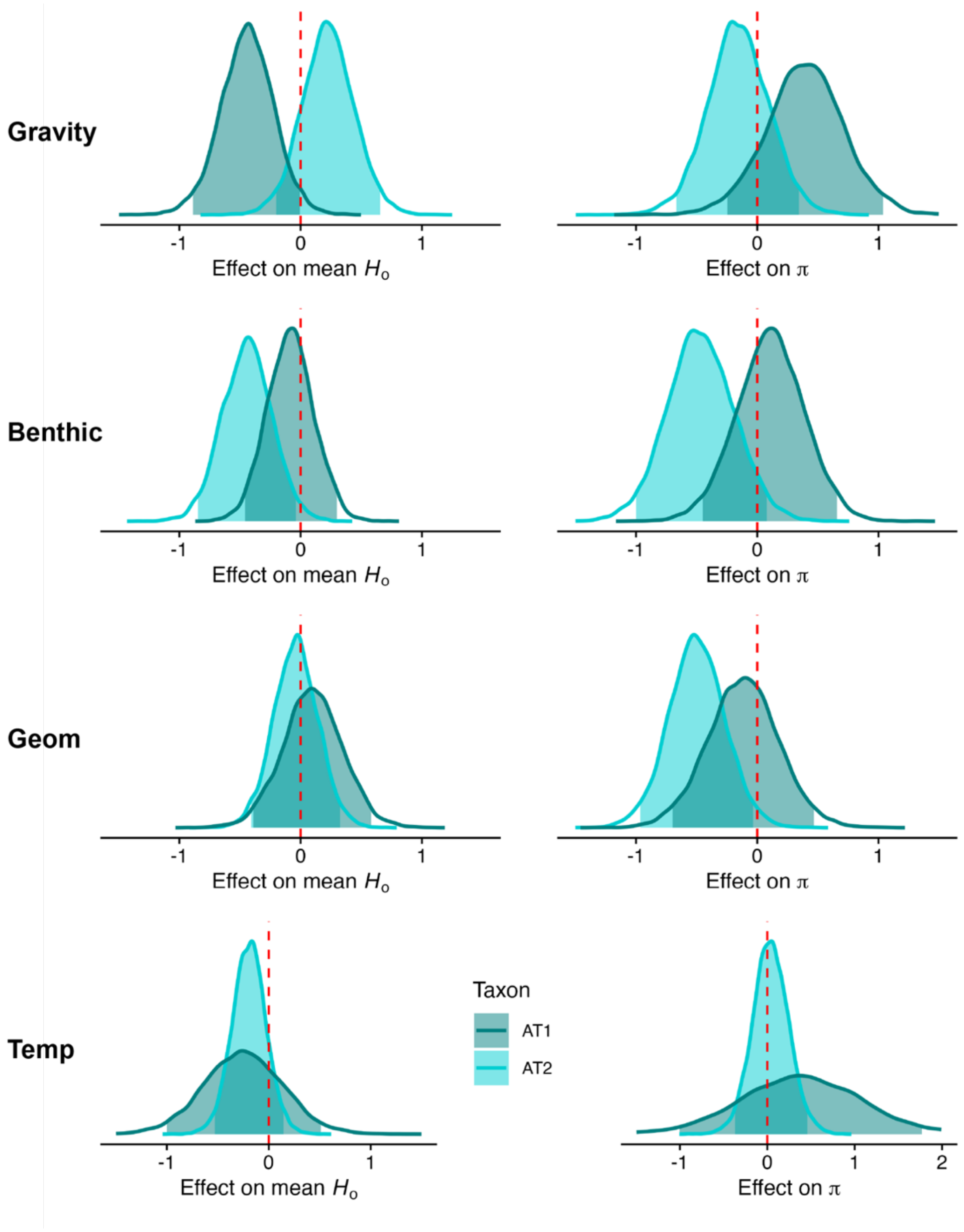


**Figure S17** Comparison of brms and INLA models shows little variation in overall relationships between habitat heterogeneity and combined or taxon-specific genetic diversity (except for gravity, which shows opposite directions, albeit not significant effects based on a 95% credible interval threshold). The INLA versions of the models shown in the main text are presented here for each of our four focal predictors, where spatial autocorrelation is incorporated using a Stochastic Partial Differential Equation (SPDE) approach.

2.7 Bayesian GLMMs extended tables

**Table S11** Results from Bayesian Linear Mixed Models performed using brms. These results correspond to the posterior distributions shown in Figure 5 in the main text. Estimates in bold are associated with confidence intervals that do not include zero, providing confidence in the strength of those estimates.

| *Focal predictor variable* | *Model* | *Coefficient estimate* | *95% credible interval* | *Significant* |
|---|---|---|---|---|
| Gravity | ho ~ taxon + gravity + (1 \| siteID) | -0.068 (0.105) | [-0.275, 0.143] | No |
| Benthic heterogeneity | ho ~ taxon + benthic + wave period + (1 \| siteID) | -0.315 (0.101) | [-0.517, -0.112] | Yes |
| Geomorphic heterogeneity | ho ~ taxon + benthic + wave period + (1 \| siteID) | -0.048 (0.102) | [-0.239, 0.158] | No |
| Mean temperature | ho ~ taxon + temperature + period + (1 \| siteID) | -0.215 (0.114) | [-0.442, 0.010] | No |

**Table S12** Results from Bayesian Linear Mixed Models performed using brms. These results correspond to the posterior distributions shown in Figure 6 in the main text.

| *Focal predictor variable* | *Model* | *AT1 estimate* | *AT1 95% credible interval* | *A2 estimate* | *AT2 95% credible interval* | *Significant* |
|---|---|---|---|---|---|---|
| Gravity | ho ~ taxon * gravity + (1 \| siteID) | -0.243 (0.130) | [-0.503, -0.008] | 0.142 (0.141) | [-0.132, 0.423] | AT1 |
| Benthic heterogeneity | ho ~ taxon * benthic + wave period + (1 \| siteID) | -0.216 (0.128) | [-0.467, 0.033] | -0.457 (0.149) | [-0.751, -0.168] | AT2 |
| Geomorphic heterogeneity | ho ~ taxon * benthic + wave period + (1 \| siteID) | 0.051 (0.152) | [-0.240, 0.363] | -0.120 (0.131) | [-0.372, 0.136] | none |
| Mean temperature | ho ~ taxon * temperature + period + (1 \| siteID) | -0.303 (0.241) | [-0.769, 0.176] | -0.203 (0.122) | [-0.448, 0.031] | none |

**Table S13** Results from Bayesian Linear Mixed Models performed using brms. Results correspond to the difference in posterior means between taxa as reported in Table S12.

| *Focal predictor variable* | *Model* | *Difference* | *95% credible interval* | *Significant* |
|---|---|---|---|---|
| Gravity | ho ~ taxon * gravity + (1 \| siteID) | 0.385 (0.178) | [0.033, 0.735] | Yes |
| Benthic heterogeneity | ho ~ taxon * benthic + wave period + (1 \| siteID) | -0.240 (0.183) | [-0.601, 0.117] | No |
| Geomorphic heterogeneity | ho ~ taxon * benthic + wave period + (1 \| siteID) | -0.171 (0.191) | [-0.550, 0.196] | No |
| Mean temperature | ho ~ taxon * temperature + period + (1 \| siteID) | 0.100 (0.251) | [-0.406, 0.597] | No |
